# Strip cropping shows promising increases in ground beetle community diversity compared to monocultures

**DOI:** 10.1101/2024.11.02.621655

**Authors:** Luuk Croijmans, Fogelina Cuperus, Dirk F. van Apeldoorn, Felix J.J.A. Bianchi, Walter A.H. Rossing, Erik H. Poelman

## Abstract

Global biodiversity is declining at an unprecedented rate, with agriculture as one of the major drivers. There is mounting evidence that intercropping can increase insect biodiversity while maintaining or increasing yield. Yet, intercropping is often considered impractical for mechanized farming systems. Strip cropping is a type of intercropping that is compatible with standard farm machinery and has been pioneered by Dutch farmers since 2014. Here, we present ground beetle data from four organically managed experimental farms across four years. Ground beetles are sensitive to changes in habitats and disturbances, and hold keystone positions in agroecosystem food webs. We show that strip cropping systems can enhance ground beetle biodiversity, while other studies showed that these increases have been achieved without incurring major yield loss. Strip cropped fields had on average 15% more ground beetle species and 30% more individuals than monocultural fields. The higher ground beetle richness in strip crops was explained by the merger of crop-related ground beetle communities, rather than by ground beetle species unique to strip cropping systems. The increase in field-level beetle species richness in organic agriculture through strip cropping approached increases found for other readily deployed biodiversity conservation methods, like shifting from conventional to organic agriculture (+19% - +23%). This indicates that strip cropping is a potentially useful tool supporting ground beetle biodiversity in agricultural fields without compromising food production.

## Introduction

Insects account for 80% of the animal species in the world and, therefore, the recently reported unprecedented rate of insect decline is cause for alarm about the state of biodiversity on Earth (Dirzo et al., 2014; Hallmann et al., 2017; Van Klink et al., 2020). An array of biodiversity metrics provide strong evidence for declines of especially terrestrial insects across all continents (Van Klink et al., 2020). This includes declines in abundance of both common and rare species (Hallmann et al., 2017; Pilotto et al., 2020; Seibold et al., 2019; van Klink et al., 2024) and in total species richness and changes in assembly composition (Blowes et al., 2022; Wagner et al., 2021). Insects are essential for crop production through their role in decomposition, pollination, pest control and sustaining food webs. Therefore, erosion of insect communities can have potentially devastating effects on ecosystem functioning, provision of ecosystem services and ultimately on human civilization (Dirzo et al., 2014). The main drivers of insect biodiversity decline are habitat loss due to conversion to agriculture, pollution, invasive species and climate change (Díaz et al., 2019; Müller et al., 2024; Wagner et al., 2021). Strategies for biodiversity conservation in conjunction with adequate food production require understanding of how biodiversity responds to agricultural management (Cozim-Melges et al., 2024; Mupepele et al., 2021; Saunders, 2020). Ideally, sustainable agricultural practices retain yield and enhance biodiversity, preventing the need to convert natural habitats to agricultural land to maintain food production (Tscharntke et al., 2021).

Increasing crop heterogeneity can facilitate biodiversity conservation in highly productive agricultural landscapes without compromising yield (Martin-Guay et al., 2018; Sirami et al., 2019). Crop diversification can enhance niche complementarity by creating heterogeneous habitats and increasing availability and diversity of resources (Lichtenberg et al., 2017; Tamburini et al., 2020). A promising crop diversification strategy is strip cropping, where crops are grown in alternating strips, wide enough for using standard agricultural machines yet narrow enough to facilitate ecological interactions among crops (Ditzler et al., 2021). Crops that are grown in strips may benefit from increased resource use efficiency, and the suppression of pests and diseases (Croijmans et al., 2024; Karssemeijer et al., 2024; Rakotomalala et al., 2023) without major yield compromises (Campanelli et al., 2023; Juventia and van Apeldoorn, 2024; Van Oort et al., 2020). Growing multiple crops on a field may foster a larger diversity of organisms than monocultures through greater plant species richness that cascades into richer herbivore and predator communities (Crutsinger et al., 2006; Cuperus et al., 2023). Moreover, the expected increase in available and potentially complementary niches within the agricultural field due to higher spatial diversity in crops can result in admixture of communities related to individual crops (Hummel et al., 2012), or the creation of completely new communities by enhanced richness of agriculture-related species, and/or the occurrence of species rarely found in agricultural fields (Rischen et al., 2021). So far, it is not well understood if and how insect communities respond to strip cropping across distinct crops.

Here, we present data of four years of pitfall trapping of ground beetles at several moments during the growing season in 14 crops at four organic experimental farms across the Netherlands where strip cropping was compared to monocultures. Ground beetle communities are sensitive to changes in farming practices and are frequently used to examine agricultural sustainability (Holland and Luff, 2000; Makwela et al., 2023; Turin, 2022). Furthermore, ground beetle species are important for maintaining ecological functions as they comprise scavengers and predators of (weed) seeds, detritivores (e.g., collembolas and earthworms) and herbivores (e.g., aphids and caterpillars). We first examine whether strip cropping fields have greater ground beetle activity density, species richness, evenness and diversity than monocultural fields. We also test for twelve abundant ground beetle genera whether their activity density is higher in strip cropping than in monocultures. Lastly, we evaluate whether ground beetle community changes are caused by admixture of communities, whether these assemblages promote species associated with agricultural or with natural ecosystems, and whether they contain rare species.

## Results

A total of 48,108 ground beetles belonging to 71 species were caught using pitfall traps over four years at four different organically managed experimental farms in The Netherlands: 40,153 at Almere; 3,777 at Lelystad; 1,126 at Valthermond; and 3,052 at Wageningen (Table S1, S2).

### Strip cropping enhances ground beetle richness

Strip cropping fields had on average 15% higher ground beetle taxonomic richness than monoculture fields after rarefaction to the number of samples of the least-sampled crop configuration (β_0_ = 0.151, SE = 0.044, p < 0.001; Fig. 1c,d). However, strip cropping fields did not harbour more species than monocultural fields with the highest ground beetle richness (β_0_ = - 0.008, SE = 0.037, p = 0.821; Fig. 1d). The difference in field-level taxonomic richness could not be explained by an increase in the number of ground beetle species per crop in strip cropping compared to monocultures. At crop-level, the 5% increase in ground beetle taxonomic richness in strip cropping was not statistically significant (Fig. 1f). Similarly, crop-level absolute evenness, inverse Simpson index and Shannon entropy did not differ significantly among crop configurations (Fig. 1h-j, S1c,d, S2). The effect of crop configuration on crop-level taxonomic richness was variable and was not associated with location or crop species. The effect ranged from 56% more species in potato in monoculture at Wageningen in 2022, to 136% more species in barley in strip cropping at Wageningen in 2020 (Fig. 1k).

**Fig. 1.**
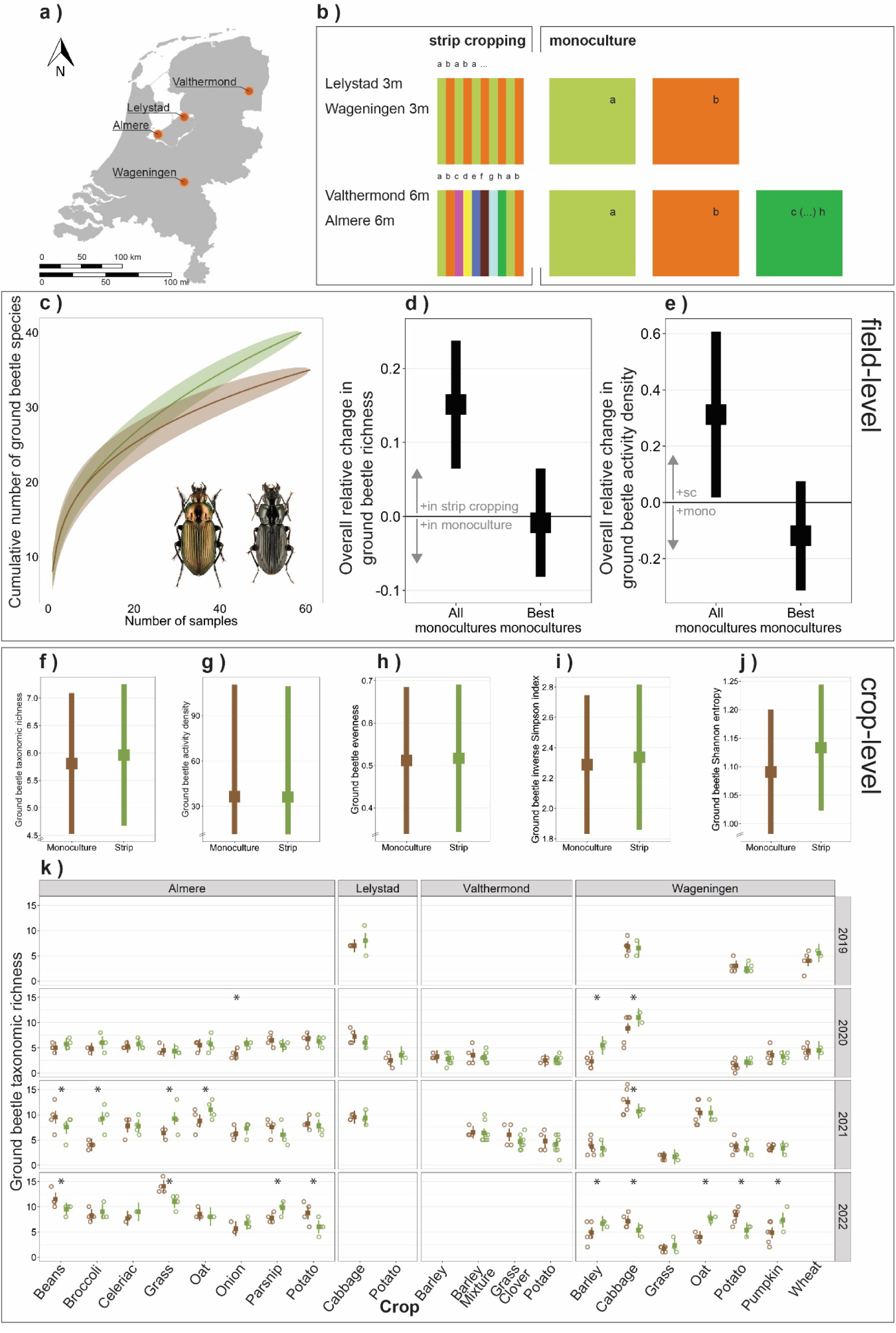
Effect of crop configuration (monoculture versus strip cropping) on ground beetle biodiversity. **(a)** Location of experimental sites in the Netherlands. **(b)** Field set-up of the two crop configurations: monoculture and strip cropping. At Lelystad and Wageningen, strip cropping consisted of 3-m wide crop strips of two crops (pairs), and multiple crop pairs were assessed. At Almere and Valthermond, strip cropping consisted of 6-m wide crop strips of eight crops combined (Fig. S10). **(c)** Sample-based species accumulation curves of all year series from monocultures (brown) and strip cropping (green), in Almere from 2021 and 2022. This was the only location where an equal number of samples were taken in the monocultures and in strip cropping on a similar area. Ground beetle species include *Poecilus cupreus* (left) and *Pterostichus melanarius* (right). Photo credit: Ortwin Bleich, retrieved from: www.eurocarabidae.de. **(d-e)** Overall relative change in field-level ground beetle **(d)** taxonomic richness and **(e)** activity density. Positive values indicate higher richness or activity density in strip cropping, negative values in monocultures. **(f-j)** Overall relative effect of crop configuration on ground beetle **(f)** taxonomic richness, **(g)** activity density, **(h)** absolute evenness, **(i)** inverse Simpson index, and **(j)** Shannon entropy. **(k)** Effect of crop configuration on ground beetle taxonomic richness for each combination of location, year and crop. Barley-mixture consists of a mixture of barley-bean (2020) or barley-pea (2021). Squares indicate estimated means, the bar indicates the 95% confidence interval. Asterisks indicate significant differences among the crop configurations. Empty panels indicate combinations of years and locations that were not sampled. When no estimated mean and confidence interval are shown, crops were not grown or sampled in that year. Open circles indicate individual year series to visualize sample size (Table S7).

### Strip cropping enhances ground beetle activity density

Ground beetle activity density was on average 30% higher in strip cropping fields than in monoculture fields (β_0_ = 0.303, SE = 0.121, p = 0.012; Fig. 1e), based on rarefaction. However, there was no significant difference in activity density between the strip cropping fields and monocultures that harboured the richest beetle communities (β_0_ = -0.110, SE = 0.088, p = 0.215). Crop-level activity density of ground beetles was not affected by crop configuration (Fig. 1g, S1a,b).

### Crop configuration alters abundance of abundant genera

We tested how crop configuration affected the abundance of twelve abundant ground beetle genera (Table S3). We analysed this separately per location as some genera only occurred at specific locations. Four genera were more abundant in strip cropped fields in at least one location (*Anchomenus, Bembidion, Harpalus* and *Nebria*), whereas four genera were more abundant in monocultures in at least one location (*Amara, Calathus, Pterostichus* and *Trechus*) (Fig 2). The other four common genera (*Blemus, Clivina, Loricera* and *Poecilus*) were not significantly influenced by crop configuration. Furthermore, no ground beetle genus showed significantly contrasting responses to crop configuration among locations.

**Fig. 2.**
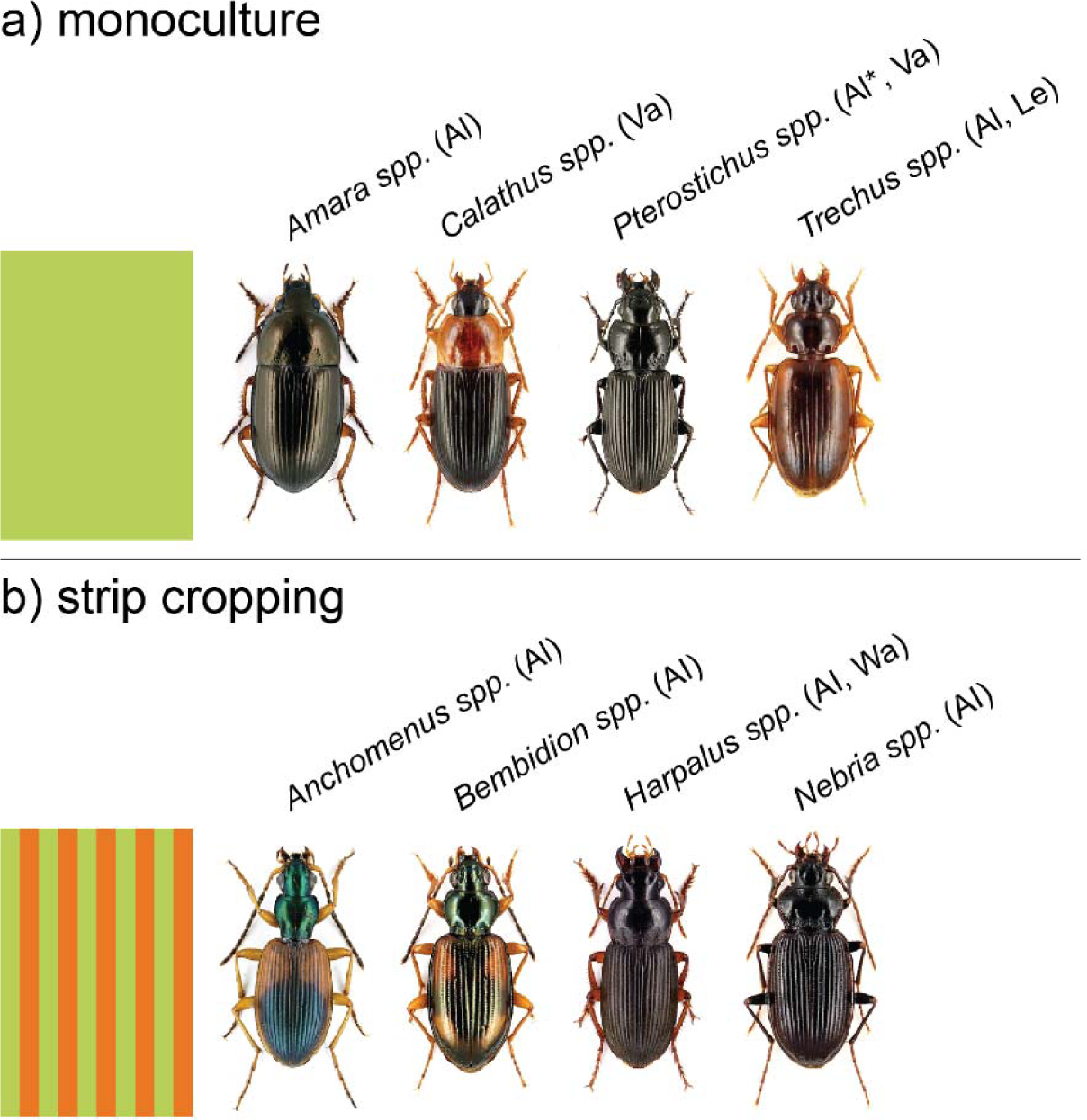
Ground beetle species associated with crop configuration (monoculture (a) versus strip cropping (b)). Results obtained by generalized linear mixed models on the twelve most common genera of ground beetles with the four locations analysed separately (Table S5). Only those genera are given for which cropping system significantly influence activity density within at least one location (α = 0.05). Locations were the genus had a higher activity density in one of the cropping systems are indicated between brackets (Al=Almere; Le = Lelystad; Va = Valthermond; Wa = Wageningen). For Pterostichus, data from Almere in 2020 was analysed separately, as models did not fit elsewise. The asterisk “*” next to Pterostichus indicates that only in 2020 the difference between monoculture and strip cropping was significant. Photo credit: Ortwin Bleich, retrieved from: www.eurocarabidae.de.

### Crop configuration alters ground beetle community composition

Ground beetle communities were significantly influenced by crop configuration, but the effects were highly dependent on the specific context created by the combination of location, year and crop (Fig. S3-S8, Table 1, S4). The context-dependency of configuration effects on ground beetle communities is illustrated in redundancy analyses of ground beetle assemblages per crop combination at Wageningen in 2021 and 2022 (Fig. S9). Here, we found distinct ground beetle communities among crop configurations for pumpkin, barley and potato in 2021, and for cabbage and oat in 2022. In the other cases the difference between crop configurations was not significant (Fig. S9, Table S5). Moreover, in all crop combinations except for potato-grass in 2021, the difference in ground beetle communities between the monocultures of the constituent crops were significant, while this was never the case for ground beetle communities of crops in strip cropping (Table S5). This indicates that strip cropping might lead to overlapping crop-related communities. However, these results could be spatially autocorrelated as samples from different crops were in closer proximity of each other in strip cropping than among monocultures (Fig S10d).

**Table 1.** Effect of crop configuration on ground beetle community composition. Results from permanova analysis using Hellinger’s transformation for data from the three locations with species level data (see Table S3 for analyses per location). Crops were a nested variable within years, as these differed among years. Years were nested in locations, as the years that were studied differed among locations. Bold letters indicate significant effects (α = 0.05).

| Location | Predictor | Df | Sum<br>sq | R2 | F | P |
| --- | --- | --- | --- | --- | --- | --- |
| All locations | Crop configuration | 1 | 1.68 | 0.01 | 7.32 | **0.001** |
|  | Location | 2 | 47.7 | 0.29 | 103.8 | **0.001** |
|  | Location : Year | 6 | 10.7 | 0.06 | 7.72 | **0.001** |
|  | Location : Year :<br>Crop species | 31 | 33.3 | 0.20 | 4.67 | **0.001** |
|  | Crop configuration : Location | 2 | 0.63 | 0.00 | 1.37 | 0.149 |
|  | Crop configuration :<br>Location : Year | 6 | 2.31 | 0.01 | 1.67 | **0.012** |
|  | Crop configuration :<br>Location : Year :<br>Crop species | 31 | 14.8 | 0.09 | 2.08 | **0.001** |
|  | *Residual* | *245* | *56.3* | *0.34* |  |  |
|  | *Total* | *324* | *167.4* | *1.00* |  |  |

### Crop configuration does not increase the number of rare ground beetle species

Among the 461 year series, we only found two rare species (following waarneming.nl): one individual of *Microlestes minutulus*, and five individuals of *Harpalus signaticornis*. This latter species was found most often in a wheat monoculture, but this was likely due to a failed crop which created a very open habitat. *Harpalus signaticornis* is known to inhabit recently disturbed, dry, open habitats with limited vegetation (Turin et al., 2012), a situation similar to this sparsely-covered monoculture. All other species were common or relatively common species (Table S2). Furthermore, most ground beetle species were either ruderal habitat specialists or eurytopic species that occur in many different habitats (Table S2).

## Discussion

It is well established that different crop types have distinct ground beetle communities (Eyre et al., 2009; Holland and Luff, 2000) and that increasing habitat diversity by including multiple crops in a field can enhance ground beetle diversity (Cuperus et al., 2024; Puliga et al., 2023). Our study shows that strip cropping increased field-level ground beetle richness by 15%. However, ground beetle communities within the same crop in a strip or in monoculture were mostly similar. This indicates that the 15% increase in richness at the field level can be mostly attributed by the higher number of crops in strip cropped fields that harboured crop-related ground beetle communities, and that there was only limited mixing of ground beetle species among crops in strip crops. This is in line with earlier findings that ground beetle movement is reduced by crop edges (Allema et al., 2014; Anderson et al., 2024). Further research on movement behaviours of ground beetles at crop edges might help explain how ground beetles distribute themselves within a strip cropping field, and whether they utilize the different resources provided by a more diverse cropping system.

The 30% higher ground beetle activity density in strip cropping fields compared to monoculture fields may be explained by a more stable and diverse habitat with refuges and alternative resources in strip crops (Ratnadass et al., 2012). Crop diversification enhances prey biomass for ground beetles (Lichtenberg et al., 2017) and increases weed seed richness as compared to monocultures (Ditzler et al., 2023), both of which reduce bottom-up control by increased food provision (Carbonne et al., 2022). Alternatively, the increase in activity density might be caused by higher movement of ground beetles, which in turn can be the consequence of food starvation (Wallin and Ekbom, 1994). Indeed, several papers show reduced abundances of herbivorous insects in strip- and intercropping (Alarcón Segura et al., 2022; Cuperus et al., 2023; Rakotomalala et al., 2023), which may be potential prey for ground beetles (Turin, 2000). Furthermore, strip cropping systems may support high herbivore reproduction in combination with a high predation pressure resulting in herbivore populations dominated by early life stages (Karssemeijer et al., 2024). Therefore, strip cropping may favour ground beetles that predate on, for instance, herbivore eggs, such as *Bembidion spp.* and *Anchomenus dorsalis*, which were more abundant in strip cropping (Finch, 1996), and disadvantaging ground beetles that feed on larger prey, such as *Pterostichus melanarius,* which was less abundant in strip cropping. Multi-taxa evaluation of future strip cropping studies will be a valuable approach to increase understanding of biomass flows and trophic interactions within diverse agricultural systems.

The 15% increase in ground beetle species richness through strip cropping is mostly caused by an increase in species (relatively) common to agricultural fields, rather than an increase in rare species or species otherwise common in other habitat types. A keystone species here might be *Pterostichus melanarius*, the most common *Pterostichus* species in our samples. *Pterostichus melanarius* can be very dominant in pitfall traps, as we regularly found hundreds of individuals per trap. This relatively large ground beetle might compete with or predate on other ground beetle species (Roubinet et al., 2018). *Pterostichus melanarius* is especially well-adapted to highly intensive agriculture (Turin, 2022, 2000), which explains why it was more abundant in monocultures previously (Ditzler et al., 2021). In turn, the reduced *P. melanarius* populations in strip cropping might allow other species to persist, such as *Harpalus spp.* and *Anchomenus dorsalis*. Therefore, strip cropping might facilitate ground beetle biodiversity by reducing the dominance of a common, competitive species.

Biodiversity gains in our study did in most cases not coincide with productivity losses (Table S6). Earlier studies on crop yield in the strip cropping fields in Lelystad and Wageningen during the years of pitfall trapping show a yield decrease when strip cropping cabbage (Carrillo-Reche et al., 2023) and wheat (Ditzler et al., 2023), whereas potato yield was unaffected by crop configuration (Ditzler et al., 2023). In Almere bean and parsnip yields were higher in strip cropping, whereas oat and onion yield were lower (Juventia and van Apeldoorn, 2024). In Valthermond crop productivity was similar for monocultures and strip cropping (Table S6). Therefore, biodiversity gains do not compromise agricultural production.

We show that changing crop configuration from monoculture to strip cropping, on average, enhances ground beetle richness by 15% and activity density by 30% within agricultural fields. These results show that strip cropping can lead to increases in biodiversity that approach those achieved by shifting from conventional to organic farming practices (+19% richness, Lichtenberg et al., 2017; +23% richness, Gong et al., 2022) and by other in-field diversification measures, like hedgerows and flower strips (+23% richness, Lichtenberg et al., 2017; +24% richness, Beillouin et al., 2021). While organic management or in-field diversification measures generally lead to lower productivity (Gong et al., 2022), strip cropping can maintain crop productivity without taking land out of production (Campanelli et al., 2023; Juventia and van Apeldoorn, 2024; Van Oort et al., 2020), although occasional yield reductions have been reported (Carrillo-Reche et al., 2023; Ditzler et al., 2023). The biodiversity gain in our study was achieved without additional crop-level diversification strategies designed for biodiversity conservation, such as cover cropping or (flowering) companion plants. This increase in biodiversity stacks on top of already higher biodiversity achieved through organic management (Gong et al., 2022; Lichtenberg et al., 2017). Furthermore, the estimated richness effects of strip cropping, i.e., no change at crop level and 15% increase at field level, are conservative because they only consider the pairwise comparison of biodiversity of two or three crops in a strip cropping configuration to monocultures, whereas the inclusion of more crops within strip cropping could further enhance ground beetle biodiversity. Also, the effect of strip cropping on ground beetle biodiversity might be more pronounced in conventional agriculture and at larger field sizes, as the relatively small-scale, organic fields that we studied might have already had a relatively high ground beetle biodiversity (Tscharntke et al., 2021). The ground beetle communities in our study were dominated by farmland and eurytopic species and contained only two rare species (Turin, 2022). Future research could test whether the inclusion of other in-field diversification measures within strip-cropped fields, such as the establishment of perennial semi-natural habitats (Rischen et al., 2021; Sirami et al., 2019) or uptake at larger spatial extents (Tscharntke et al., 2021) would allow ground beetles with other habitat preferences to establish in agricultural fields.

## Materials and methods

### Study area

A multi-location study was conducted on four organic farms across the Netherlands (Fig. 1a). Three experimental farms were managed by Wageningen University & Research (Lelystad, Valthermond, Wageningen) and one commercial farm was managed by *Exploitatie Reservegronden Flevoland* (ERF B.V.) located in Almere. All four locations contained both strip cropping and monocultural crop fields, but differed in soil type, establishment year of the strip cropping experiment, number of crops grown, length of the crop rotation, number of sampled crops and sampling years, and farm and landscape characteristics such as percentage of on-farm semi-natural habitat (SNH), mean field size, and landscape configuration (Fig. 1b, S10). The locations Almere and Lelystad were located in a homogeneous, open polder landscape characterized by intensive arable crop production and non-crop habitats consisting of grass margins, tree lines and watercourses. Valthermond was located in an open, reclaimed peat landscape with intensive arable crop production characterized by long and narrow fields separated by grassy margins and ditches and limited areas of woody elements. The site at Wageningen was located in a more complex landscape with smaller field sizes, and non-crop habitat consisting of woodlots, hedgerows, tree lines, ditches and farmyards.

### Experimental lay-out

At Almere and Valthermond the crops in strip cropping were all grown alongside each other, whereas at Lelystad and Wageningen two alternating crops (crop pairs) were grown alongside each other (Fig. 1b, S10). At each location, strip cropping and monoculture fields were always paired on the same experimental field (Fig. S10). At Almere eight different crops were grown in alternating strips of 6 m width, including celeriac (*Apium graveolens* var. *rapaceum*), broccoli (*Brassica oleracea* var. *italic*), oat (*Avena sativa*), onion (*Allium cepa*), parsnip (*Pastinaca sativa*), faba bean (*Vicia faba*), potato (*Solanum tuberosum*) and a mix of ryegrass and white clover referred to as grass-clover (*Lolium perenne/Trifolium repens*) (Fig. S10a). At Lelystad four different crop pairs were grown in alternating strips of 3 m width, including carrot (*Daucus carota subsp. sativus)* and onion, white cabbage (*Brassica oleracea var. capitata*) and wheat (*Triticum aestivum*), sugar beet (*Beta vulgaris*) and barley (*Hordeum vulgare)*, and potato and ryegrass (Fig. S10b). At Valthermond eight different crops were grown in alternating strips of 6 m width, including potato, barley, barley mixed with broad bean (*Vicia faba*) in 2020, barley mixed with pea (*Pisum sativum*) in 2021, sweet corn (*Zea mays convar. saccharata var. rugosa*), sugar beet, common bean (*Phaseolus vulgaris*), and grass-clover (Fig. S10c). At Wageningen three different crop pairs were grown in alternating strips of 3 m width, including white cabbage and wheat (2019 – 2021) or oat (2022), barley and pumpkin (*Cucurbita maxima*, 2020 - 2022) or bare soil due to crop failure (2019), and potato and ryegrass (Fig. S10d). The crop combinations and neighbors were selected based on literature, expert knowledge and experience of functionality in terms of expected advantages for yield and pest and disease control. Large scale monoculture plots (0.25 ha to 2.30 ha) served as reference, hereafter referred to as monoculture (Fig. S10). At Lelystad, Valthermond and Wageningen not each crop grown in strips was present as monoculture in each year, but only those crops for which a monoculture was present were sampled (Fig. 1k, Table S7). All fields were managed according to organic regulations, yet at each location fertilization and weed management reflected regional practices and were adjusted to local soil conditions. Flower strips were sown within the experimental fields at Almere and Valthermond (Fig. S10, Table S8).

### Sampling

The ground beetle community was sampled using pitfall traps in all crops for which both a monoculture and strip cropping field were present, at each location and in multiple rounds per year between March and September. The sampled crops, number of rounds and number of pitfalls differed per year and per location (Table S7). Pitfall traps consisted of a transparent plastic cup (9.2 cm diameter, 14 cm height) placed in the soil so the top of the cup was flush with the soil surface. We did not use funnels inside the cups. Pitfalls were filled with approximately 100 ml water mixed with non-perfumed soap (around 3 cm heigh water level) and covered with a black plastic rain guard (12.5 cm diameter) around 2 to 5 cm above the soil surface. Pitfall traps were placed in the center of a strip (1.5 – 3 meters from the edges of the strip), and at least 10 meters from field edges at fixed locations at different moments in a year. Furthermore, traps were mostly placed at equal distances from the field edges in strip-cropped and monocultural fields. At Almere and Valthermond, the number of pitfalls in monocultures and strip cropping was the same. At Wageningen and Lelystad, the number of pitfalls in strip cropped fields was usually lower than in the monoculture because the area of the strips was only half of that of the monocultures.

### Ground beetle identification

Ground beetles were identified up to species level for Almere in 2021 and 2022, Lelystad and Wageningen. At Almere in 2020, identification was kept at the genus level due to an extremely high abundance of *Pterostichus melanarius/niger* and *Poecilus cupreus/versicolor*, and the associated high time investment for identification. At Valthermond, due to labour constraints, we chose to consider species that are complicated to distinguish only up to genus level (like *Poecilus cupreus/versicolor*, *Harpalus spp.* other than *H. rufipes* and *H. affinis*, *Bembidion spp.*). For all analyses, year series were made, in which all ground beetle catches from the same pitfall trap were pooled per year. We examined habitat preferences of ground beetle using Turin et al. (2022) and ground beetle rarity using www.waarneming.nl (checked on 15-01-2025).

### Statistical analyses

We used R, version 4.2.2 for all statistical analyses.

#### Effect of crop configuration on field-level richness and activity density

To analyse the difference in species richness and activity density between monocultures and strip cropping configurations at field level, we used rarefaction of samples within the same field. To rarify to an equal sampling intensity, we calculated the average cumulative number of species or individuals within x year series, where x is the largest number of year series available for the crop configuration comparison (correcting for unequal sampling between monocultures and strip cropping, or for missing samples). Next, we calculated the relative change due to strip cropping by subtracting the number of species or individuals found in the monoculture field from the number found in the strip cropping field and then dividing the result by the number of species or individuals in the monoculture (Zou et al., 2020). This gave the relative change centered around zero, where negative values indicated higher richness in monocultures and positive values higher richness in strip cropping. We then analysed this data using Generalized Linear Mixed Models (GLMM) with a Gaussian distribution and assessed whether the intercept deviated significantly from zero. As random variables, we used location and year, with year nested in location. We ran these analyses using a dataset that included all comparisons among monocultural fields and strip cropping fields of all locations. However, as in this case the strip cropped field was compared with both monocultural fields of the corresponding crop pair, we also tested the effect of field-level richness using only one monoculture per strip cropped field. To obtain a conservative estimate of the effect of strip cropping, we chose the monocultural fields with the highest taxonomic richness or activity density among the constitutive crops of the strip cropping fields. Generalized linear models were run using the glmmTMB package (Brooks et al., 2017) and tested for model fit using the DHARMa package (Hartig, 2018).

#### Effect of crop configuration on crop-level biodiversity

To quantify biodiversity we used five variables: (1) activity density, the total number of ground beetles found per year series; (2) taxonomic richness, the total number of species or genera (lowest taxonomic level available) found per year series; (3) the inverse Simpson index, the inverse of the sum of proportions of different species over the total abundance (Simpson, 1949); (4) absolute evenness, the number of effective species calculated by dividing the inverse Simpson index by the taxonomic richness (Williams, 1964); and (5) Shannon entropy (Shannon and Weaver, 1949). We chose absolute evenness as our measure for evenness, as this method removes the richness component from the inverse Simpson index and adheres to all requirements for an evenness index (Smith and Wilson, 1996; Tuomisto, 2012). We included both the inverse Simpson index and Shannon entropy as the former is more sensitive to changes in evenness and the latter to species richness (DeJong, 1975).

To analyse the effect of crop configuration on total ground beetle activity density, taxonomic richness, evenness, inverse Simpson index and Shannon entropy we used GLMM. We constructed models for each response variable, using data from all four locations. In these models we included crop configuration (monoculture or strip cropping) as a fixed factor. We included location, year and crop as nested random variables in these models, with crop nested in year and year in location. To quantify and visualize the variation in responses between locations, years and crops, we ran generalized linear models (GLM) with a variable that combined these three variables into one, which was also included as a fixed factor. Here, we also included the interaction between crop configuration and the combined variable for crop, location and year. For the model on activity density we used the negative binomial distribution (log link function), as this was count data; for richness, evenness and Shannon entropy we used Gaussian distribution; and for the inverse Simpson index we used a gamma distribution (inverse link function) as a Gaussian distribution did not fit well. We used the ‘DHARMa’ package to validate model assumptions (Hartig, 2018). We used estimated marginal means to assess differences between monoculture and strip cropping (Hothorn et al., 2016; Lenth, 2018).

To analyse the effect of crop configuration on the activity density of the twelve most common ground beetle genera, we used another GLMM with negative binomial distribution. Fixed effects were crop configuration, location and their interactions, and crop nested in year was added as a random effect. As certain genera only occurred at specific locations, we only included locations where the genus was found. All models were tested for model fit using the DHARMa package. We used estimated marginal means to assess differences between monoculture and strip cropping.

#### Community composition of crop configurations and crops

To assess whether crop configurations have distinct ground beetle communities we used permanova with a Hellinger transformation (with 999 permutations) using the “vegan” package (Oksanen et al., 2013). We used a Hellinger transformation to give more weight to rarer species, thus accounting for highly abundant species in some samples (such as *P. melanarius* and *H. rufipes*). We only included species that occurred in at least 3% of the pitfall samples to avoid strong influence of very rare species. We used data from locations and years where pitfall catches had been identified to species level (Almere 2021/22, Lelystad and Wageningen). We considered four models, one for all locations combined and one for each location separately. In all models we included crop configuration and the nested variables of location (whenever applicable), year and crop as fixed factors. We also analysed the interaction between crop configuration and this nested structure of location, year and crop. To visualize these results, we used non-metric multidimensional scaling (NMDS) on Hellinger transformed data. To show whether crop configurations have distinct ground beetle communities, we used redundancy analysis (RDA) on Hellinger transformed data. Here, we again conducted four analyses, one for all locations combined and one for each of the three considered locations. We only used crop configuration as a predictor to force RDA to show any change in ground beetle community associated with crop configuration. As such, only one RDA axis was created per model, which was plotted against the first principal component describing the residual variation.

Due to the large influence of location and year on ground beetle communities, visualizing any general effects of strip cropping on these communities using all available data was challenging. To address this, we conducted RDA and visualized the effect of crop configuration on a subset of the data from one location and one year. We chose the data from Wageningen in 2021 and 2022 for this analysis because it provided a set-up where strip cropping of two crops could be compared with their constituent monocultures within the same experimental fields. Here, we used both crop configuration, crop and their interaction term as explanatory variables for each crop pair separately. To analyse whether ground beetle communities significantly differed among combinations of crops and crop configurations we ran pairwise permanova on all three fields and two years separately using the “pairwiseAdonis” package (Martinez Arbizu, 2020).

## Acknowledgements

First, we are grateful to Ron Anbergen, Martine Arkema, Bart Burger, Jonas Driessen, Michiel van de Glind, Angelo Grievink, Roelof Gruppen, Rolinde de Haan, Willem Hendriks, Nashita Maniran, Marina Martino, Sara Michellin, Ciska Nienhuis, Ralph Rustom, Simone Verdonschot, Pim Vrehen, Rik Waenink, and Xiaoshen Wang for their contribution to data collection. This work could not have been completed without the occasional help of many colleagues and students, for this we thank Zhaoqi Bin, Lenora Ditzler, Hilde Faber, Merel Hofmeijer, Gabriel Joachim, Stella Juventia, Peter Karssemeijer, Nelson Ríos Hernández, and any other people that helped. Thanks go to Roy Michielsen and Dirk van de Weert, for maintaining the Almere fields; the team at Wageningen Field crops, and in particular Joost Rijk and Laurens van Run, for maintaining the Lelystad fields; Gerard Hoekzema for maintaining the Valthermond fields; Olivia Elsenpeter, Esther Hofkamp, Titouan le Noc and the Unifarm staff, and in particular Andries Siepel and Peter van der Zee, for maintaining the Wageningen fields. We thank Ortwin Bleich for allowing us to use his ground beetle pictures. We are grateful to Daan Mertens, Marcel Dicke, Liesje Mommer and Thijs Fijen for constructive feedback on earlier versions of this manuscript. Lastly, we thank editors Bernhard Schmid and Sergio Rasmann, and three anonymous reviewers for their extensive and constructive feedback.

## Statements

### Author contributions

*Luuk Croijmans:* Conceptualization, Methodology, Formal analysis, Investigation, Data curation, Writing – original draft, Writing – review & editing, Visualization, Supervision. *Fogelina Cuperus:* Conceptualization, Methodology, Formal analysis, Investigation, Data curation, Writing – original draft, Writing – review & editing, Visualization, Supervision. *Dirk F. van Apeldoorn:* Conceptualization, Writing – review & editing, Visualization, Supervision, Project administration, Funding acquisition. *Felix J.J.A. Bianchi:* Conceptualization, Writing – original draft, Writing – review & editing. *Walter A.H. Rossing:* Conceptualization, Writing – original draft, Writing – review & editing, Project administration, Funding acquisition. *Erik H. Poelman:* Conceptualization, Methodology, Writing – original draft, Writing – review & editing, Supervision, Project administration, Funding acquisition.

### Ethics approval

This research was not reviewed by an institutional or governmental regulatory body as the work was performed on invertebrates.

### Availability of data and materials

Data and scripts are publicly available from 4TU.ResearchData via the following DOI: https://doi.org/10.4121/bcf78320-aaa6-428f-acf6-2eb436baa13e (Croijmans et al., 2025).

