## Supplemental figures and tables for "Strip cropping shows promising increases in ground beetle community diversity compared to monocultures"

**Supplementary figures**

 
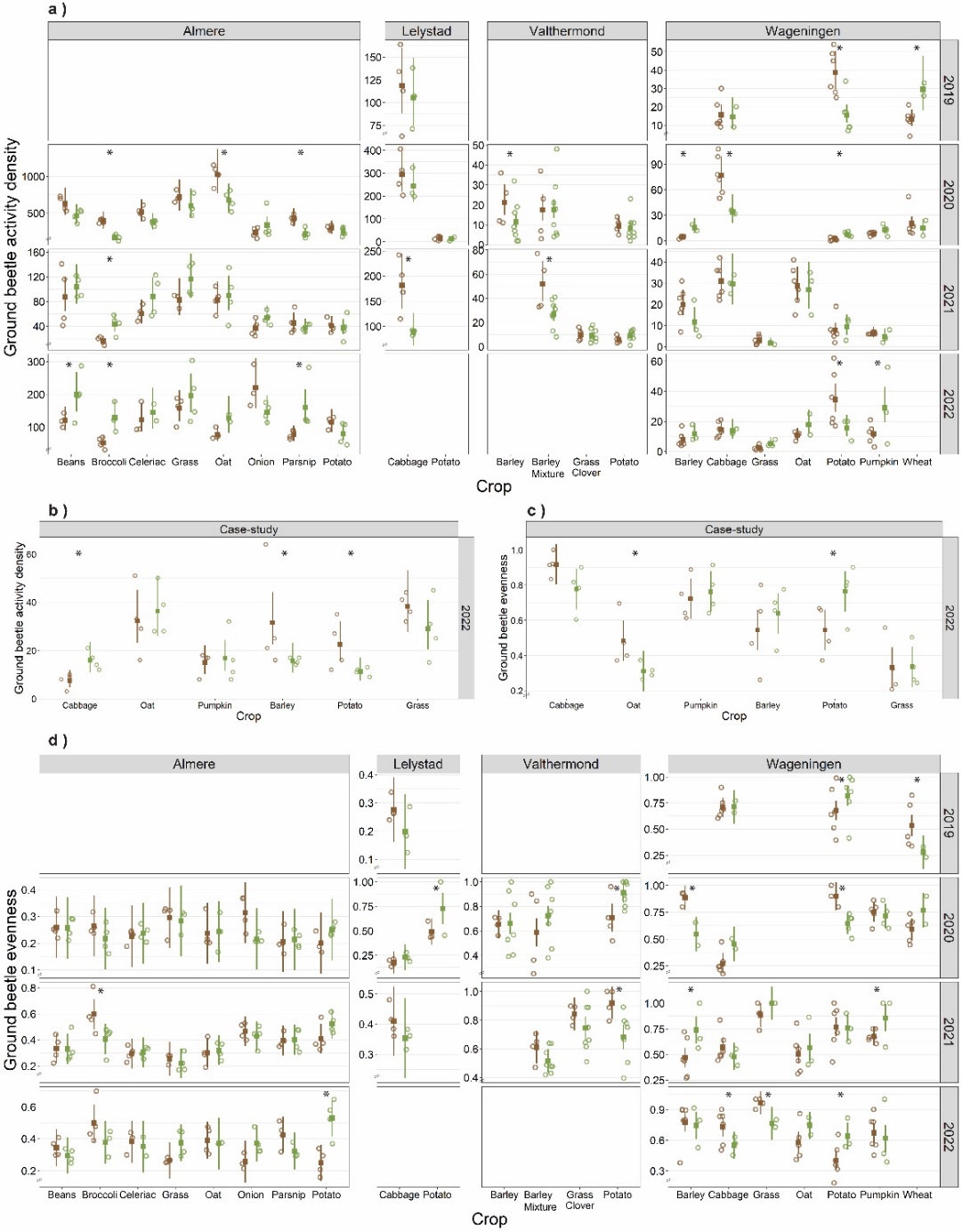

**Fig. S1. Effect of crop configuration on ground beetle activity density and absolute evenness.** Effect of crop configuration on activity density **(a)** and absolute evenness **(d)** per combination of location (columns), year (rows) and crop (x-axis); and the effect of crop configuration on activity density **(b)** and absolute evenness **(c)** in the case-study in Wageningen, consisting of two early spring sampling rounds in 2022. Activity density here is the total number of ground beetles captured in year series, which might differ in sampling frequency (Table S7). Absolute evenness is calculated from the total number of ground beetles captured in year series. Empty panels indicate combinations of years and locations that were not sampled. Squares indicate estimated means, the bar indicates the 95% confidence interval. When no estimated mean and confidence interval are shown, then those crops were not grown or sampled in that year. Asterisks indicate significant differences among the crop configurations (α = 0.05). Open circles indicate individual year series (Table S7).

  
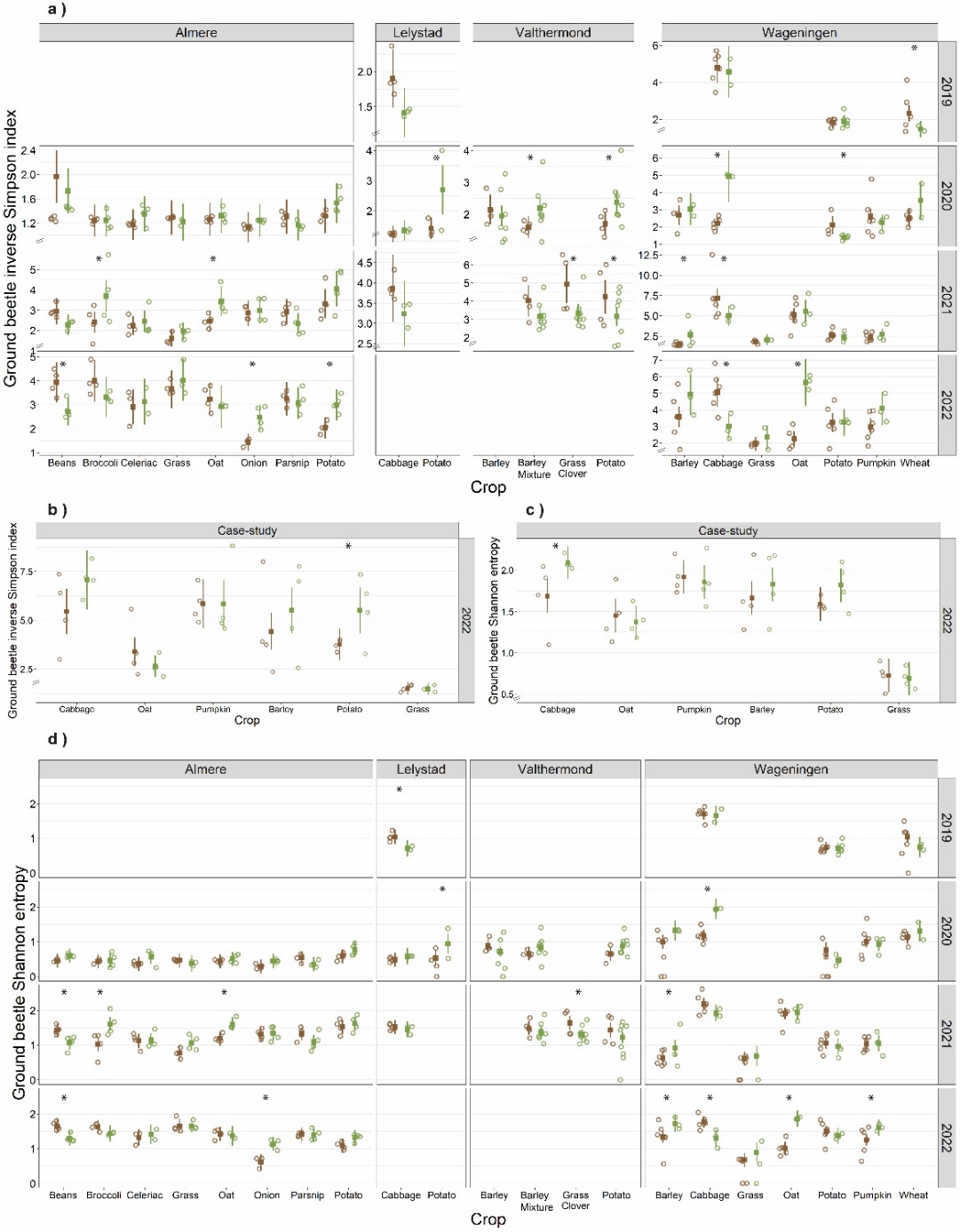

**Fig. S2. Effect of crop configuration on ground beetle inverse Simpson index and Shannon entropy.** Effect of crop configuration on inverse Simpson index **(a)** and Shannon entropy **(d)** per combination of location (columns), year (rows) and crop (x-axis); and the effect of crop configuration on inverse Simpson index **(b)** and Shannon entropy **(c)** in the case-study in Wageningen, consisting of two early spring sampling rounds in 2022. Inverse Simpson index and Shannon entropy were calculated from the total number of ground beetles captured in year series. Empty panels indicate combinations of years and locations that were not sampled. Squares indicate estimated means, the bar indicates the 95% confidence interval. When no estimated mean and confidence interval are shown, then those crops were not grown or sampled in that year. Asterisks indicate significant differences among the crop configurations (α = 0.05). Open circles indicate individual year series (Table S7).

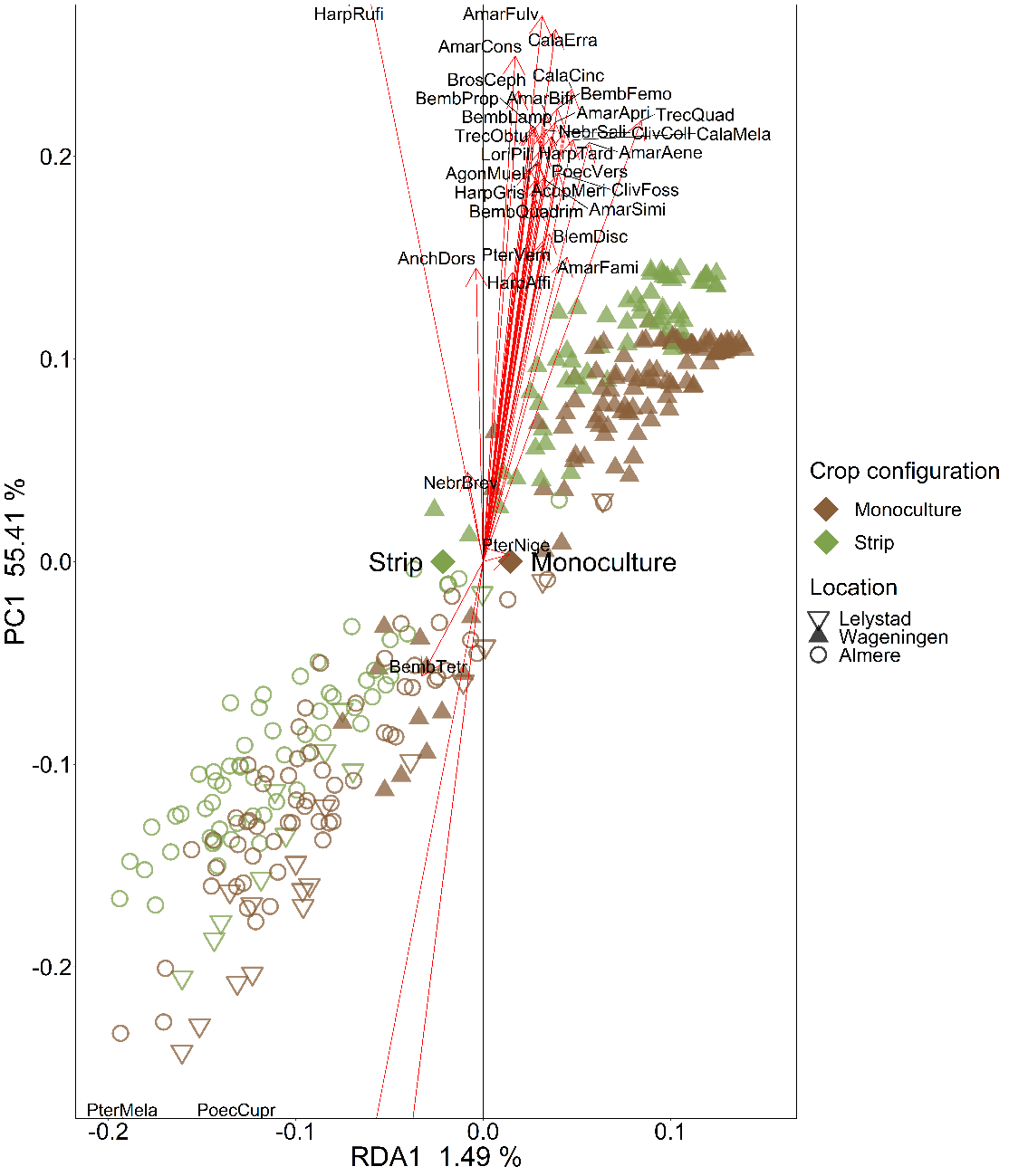

**Fig. S3. The effect of crop configuration on ground beetle community composition using redundancy analysis (RDA).** Data from all datasets that included species level data (Almere (○), Lelystad (▽), Wageningen (▲)) are used for visualizing the first RDA and PC axis. Each axis shows the percentage explained variation. Colour of the dots indicates crop configuration (brown = monoculture, green = strip cropping), and the direction of each crop configuration on the RDA axis (x-axis) is mentioned. Red arrows indicate species placement and name codes are given close to the tip of the arrow, meaning of the name codes can be found in Table S1.

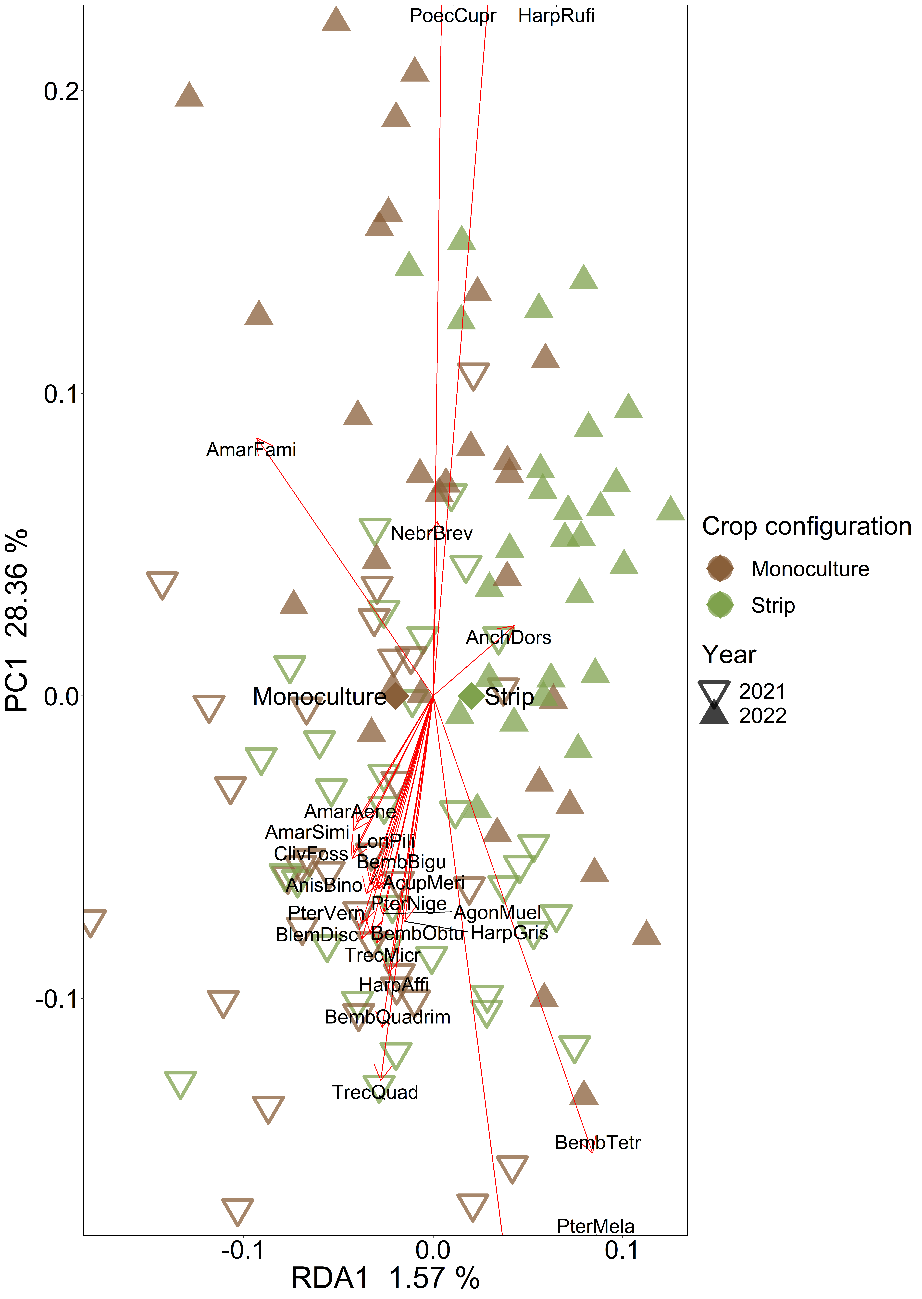

**Fig. S4. The effect of crop configuration on ground beetle community composition in Almere, using redundancy analysis (RDA).** The first RDA and PC axis are visualized. Each axis shows the percentage explained variation. Shape of the dots indicates year (▽ = 2021, ▲ = 2022), colour of the dots indicates crop configuration (brown = monoculture, green = strip cropping), and the direction of each crop configuration on the RDA axis (x-axis) is mentioned. Red arrows indicate species placement and name codes are given close to the tip of the arrow, meaning of the name codes can be found in Table S1.

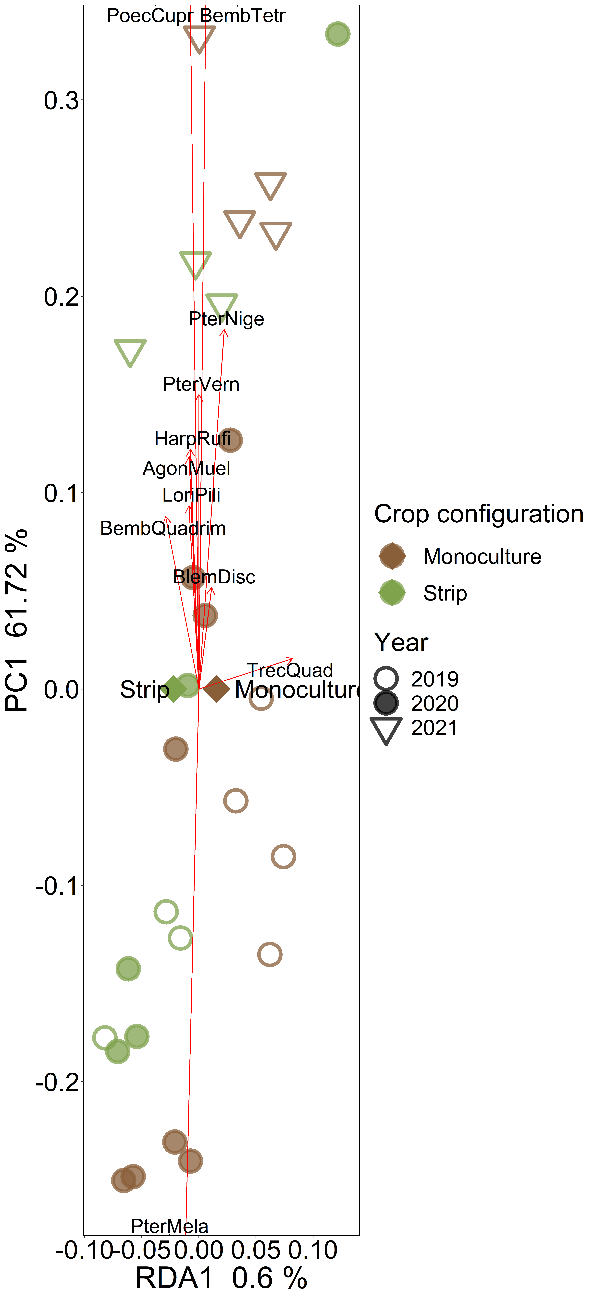

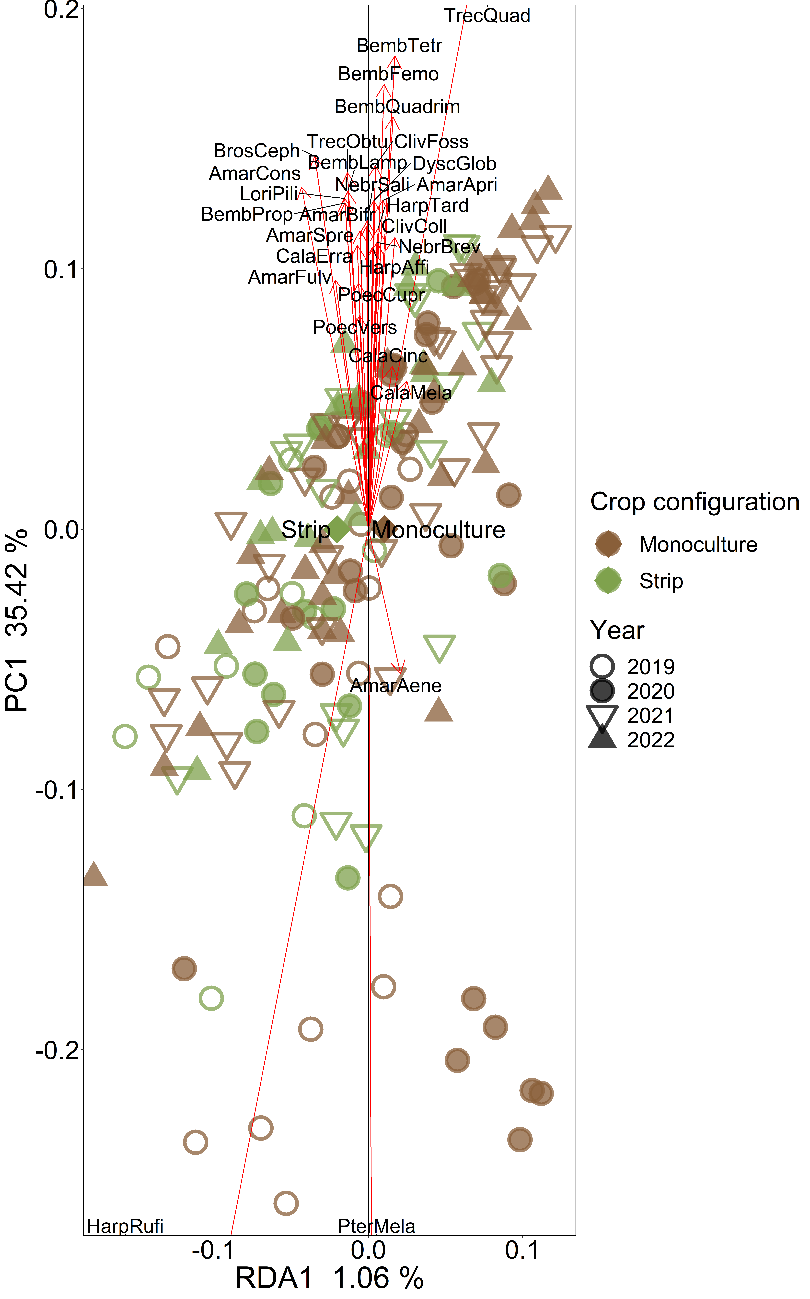

**Fig. S5. The effect of crop configuration on ground beetle community composition in Lelystad (left) and Wageningen (right) , using redundancy analysis (RDA).** The first RDA and PC axis are visualized. Each axis shows the percentage explained variation. Shape of the dots indicates year (○ = 2019, ● = 2020, ▽ = 2021, ▲ = 2022), colour of the dots indicates crop configuration (brown = monoculture, green = strip cropping), and the direction of each crop configuration on the RDA axis (x-axis) is mentioned. Red arrows indicate species placement and name codes are given close to the tip of the arrow, meaning of the name codes can be found in Table S1.

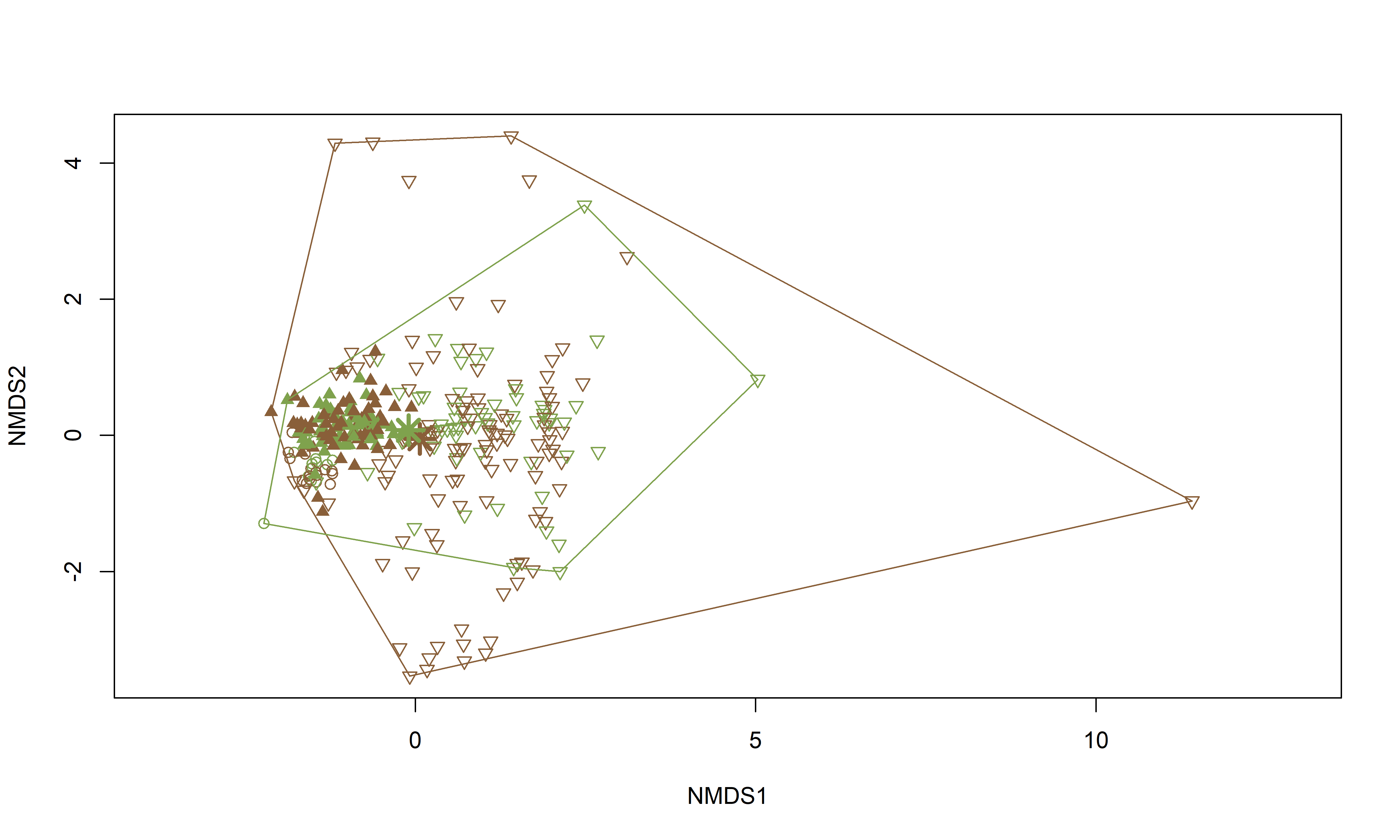

**Fig. S6. The effect of crop configuration on ground beetle community composition, using non-metric multidimensional scaling (NMDS).** Data from all datasets that included species level data (Almere (○), Lelystad (▽), Wageningen (▲)) are used for visualizing the two NMDS axes. Colour of the dots indicates crop configuration (brown = monoculture, green = strip cropping), and the centroids of each crop configuration are indicated with a large asterisk (*).

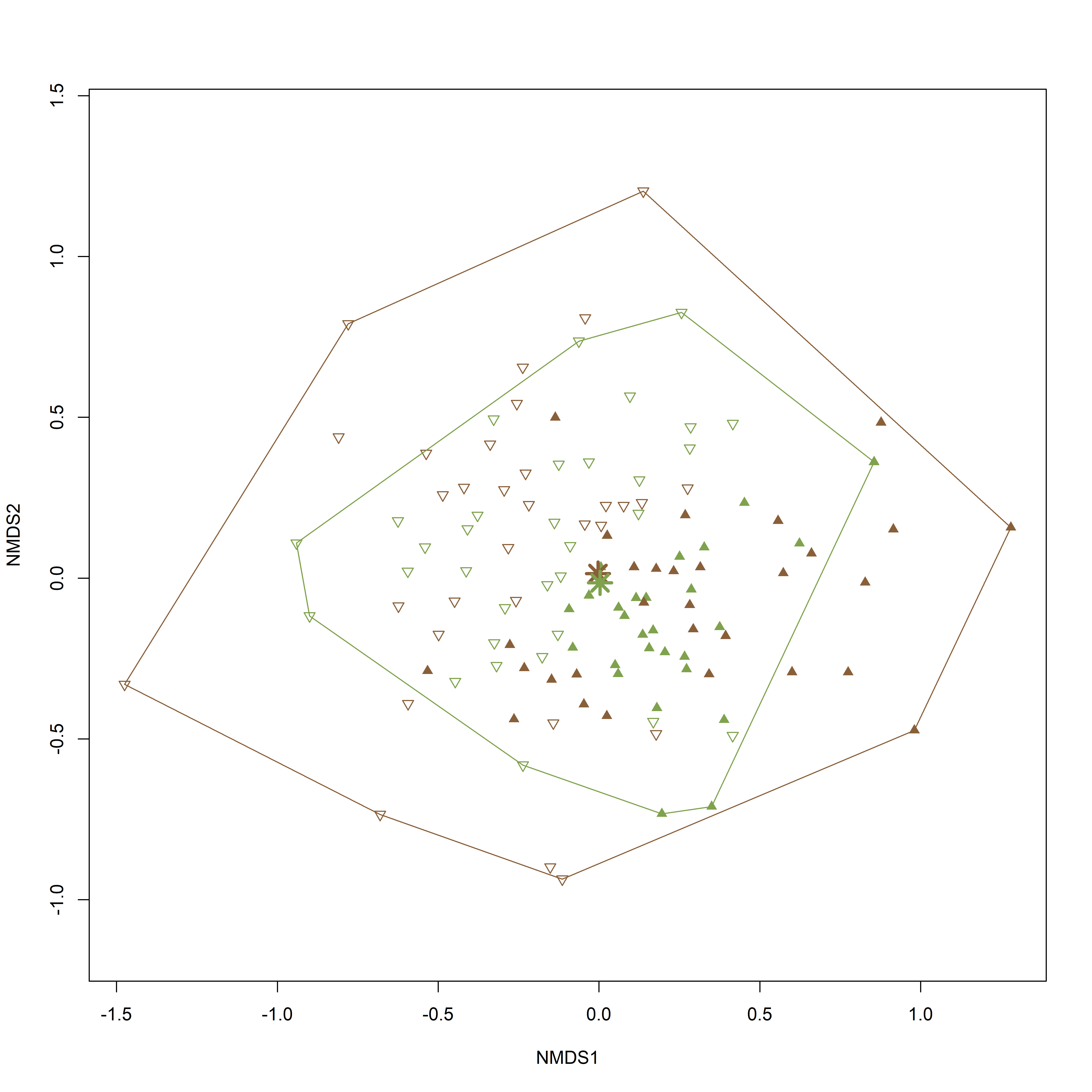

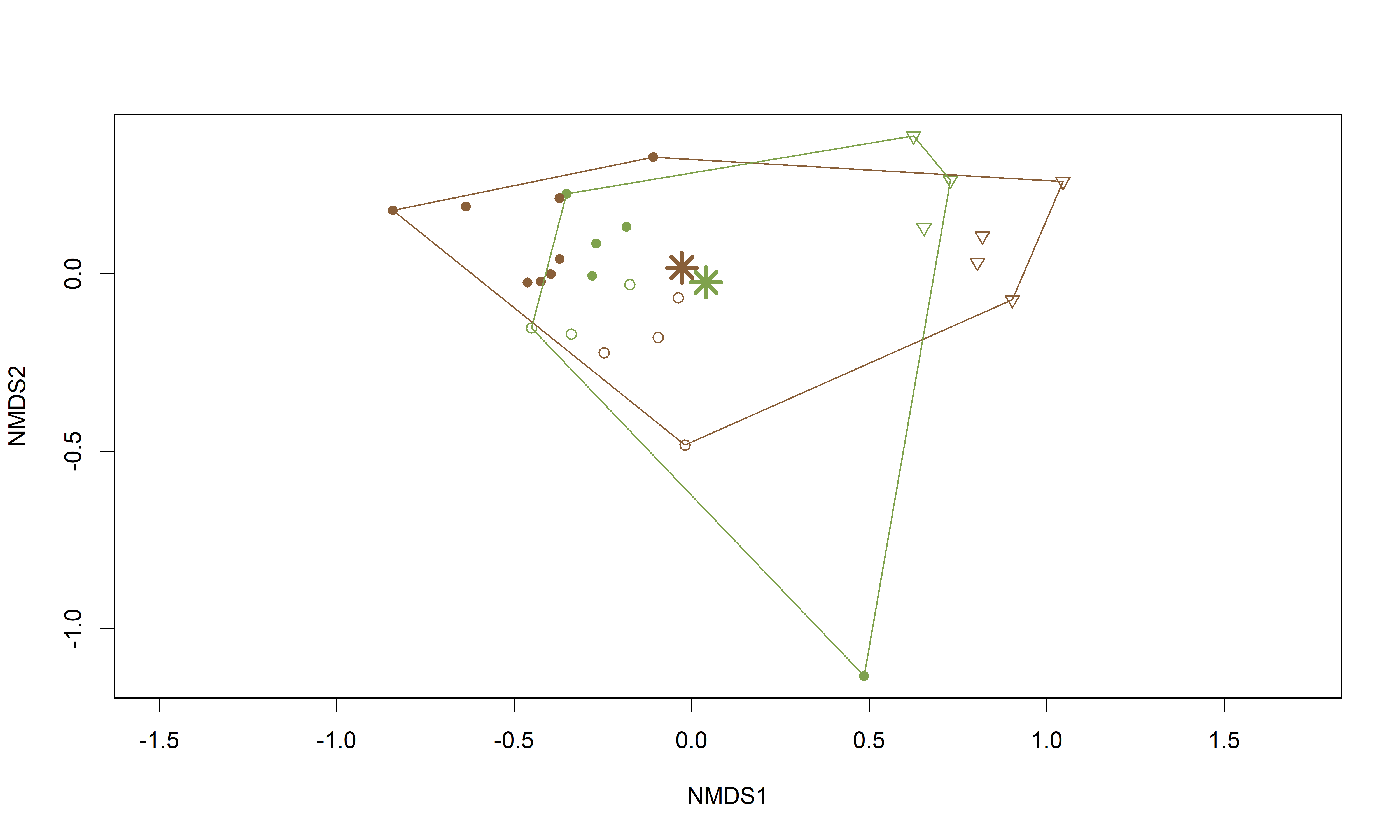

**Fig. S7. The effect of crop configuration on ground beetle community composition in Almere (top) and Lelystad (bottom) , using non-metric multidimensional scaling (NMDS).** The two NMDS axes are visualized. Shape of the points indicates year (○ = 2019, ● = 2020, ▽ = 2021, ▲ = 2022), colour of the dots indicates crop configuration (brown = monoculture, green = strip cropping), and the centroids of each crop configuration are indicated with a large asterisk (*).

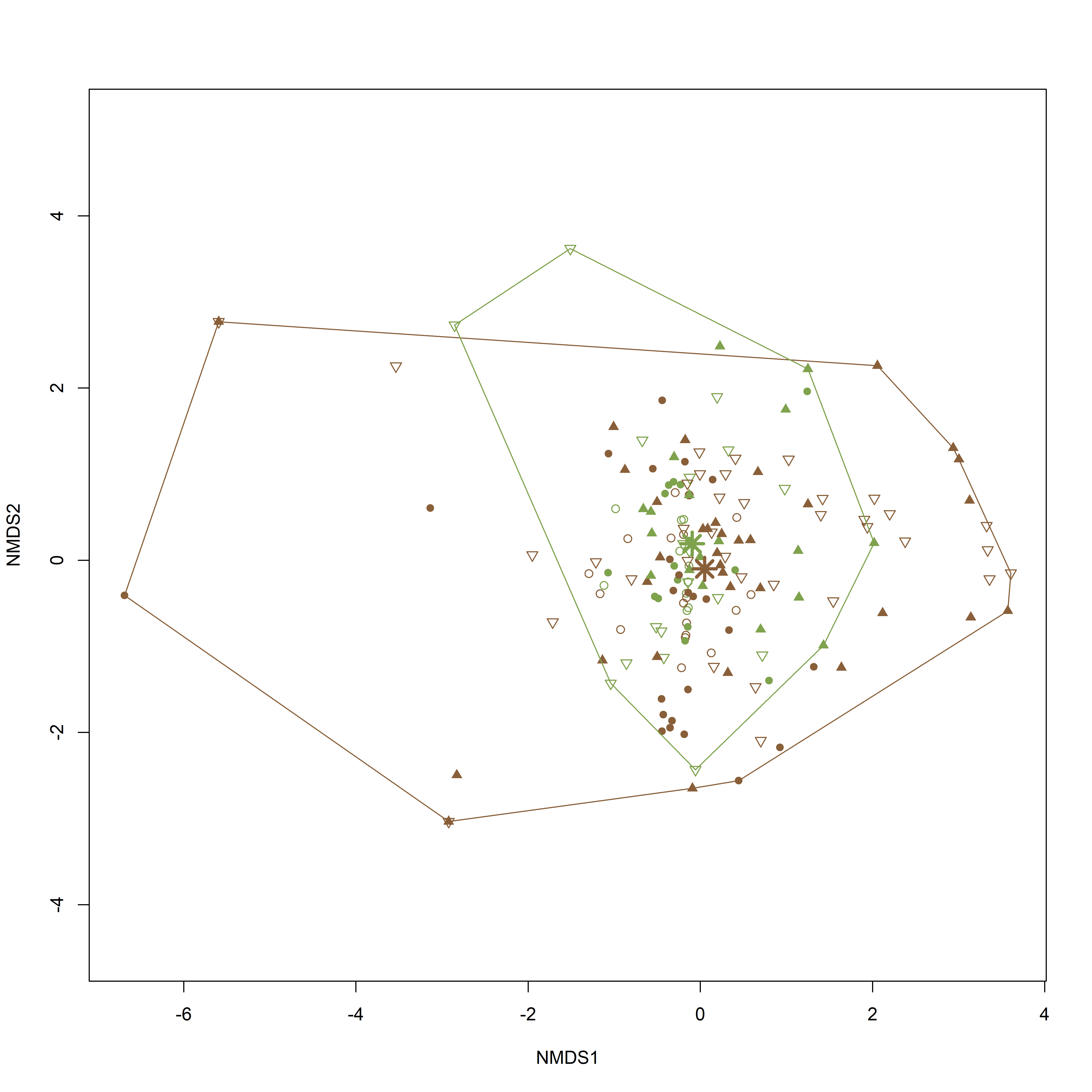

**Fig. S8. The effect of crop configuration on ground beetle community composition in Wageningen, using non-metric multidimensional scaling (NMDS).** The two NMDS axes are visualized. Shape of the points indicates year (○ = 2019, ● = 2020, ▽ = 2021, ▲ = 2022), colour of the dots indicates crop configuration (brown = monoculture, green = strip cropping), and the centroids of each crop configuration are indicated with a large asterisk (*).

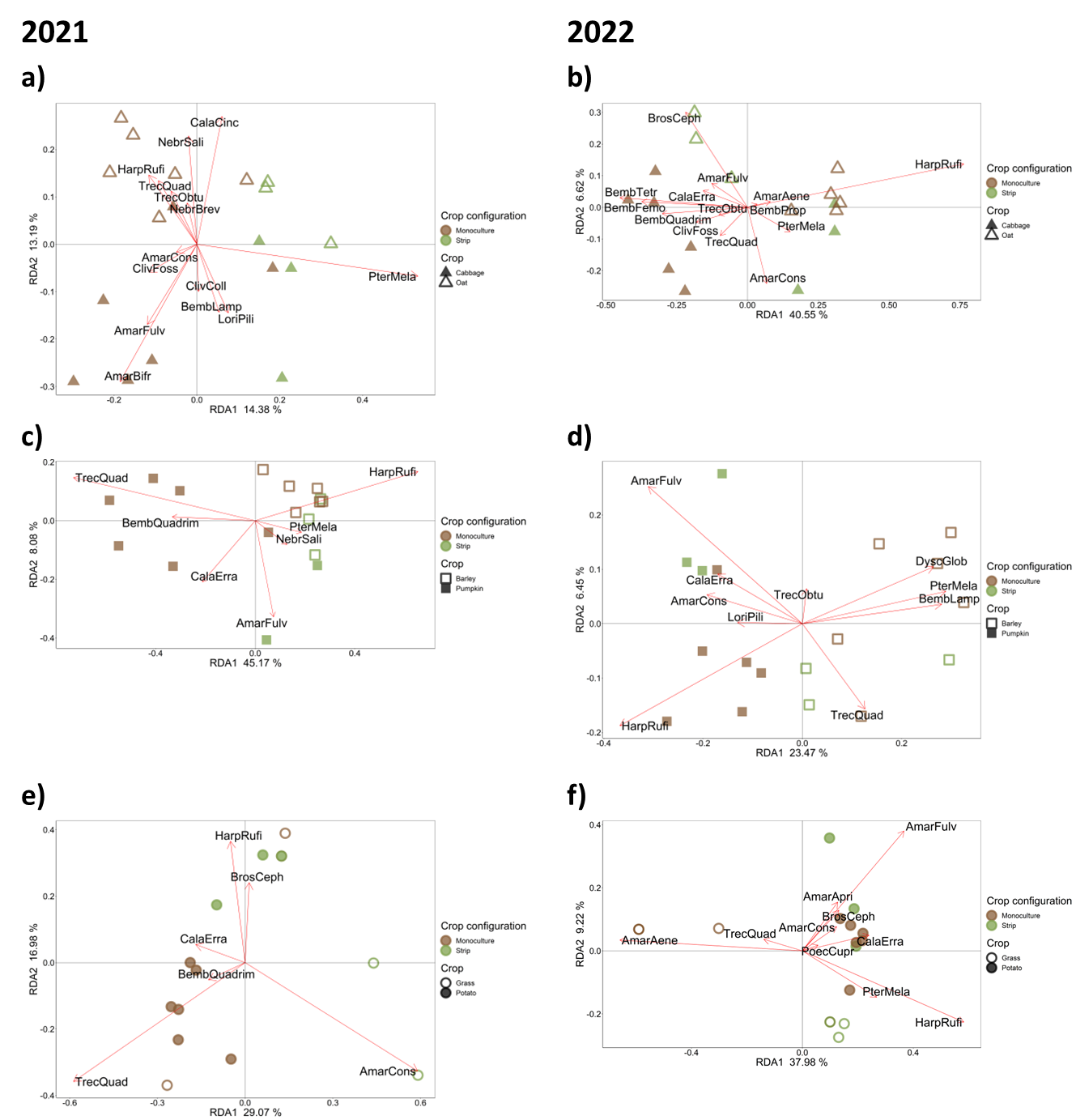

**Fig. S9. Effect of crop configuration and crop on ground beetle community composition in Wageningen.** Here, we use data from Wageningen in 2021 (left column; panels a, c, e) and 2022 (right column; panels b, d, f) from three fields including three crop pairs being cabbage and oat (panel a and b), barley and pumpkin (panel c and d), and grass and potato (panel e and f). The first and second RDA axis are visualized. Each axis shows the percentage explained variation. Colour of the dots indicates crop configuration (brown = monoculture, green = strip cropping), shapes indicates crop (□ = barley, ■ = pumpkin, ○ = grass, ● = potato, △= oat, ▲ = cabbage). The red lines and abbreviated names indicate how specific ground beetles species correlate with the RDA axes, species names are given in Table S1.

**Fig. S10. Field maps**

**A Almere**

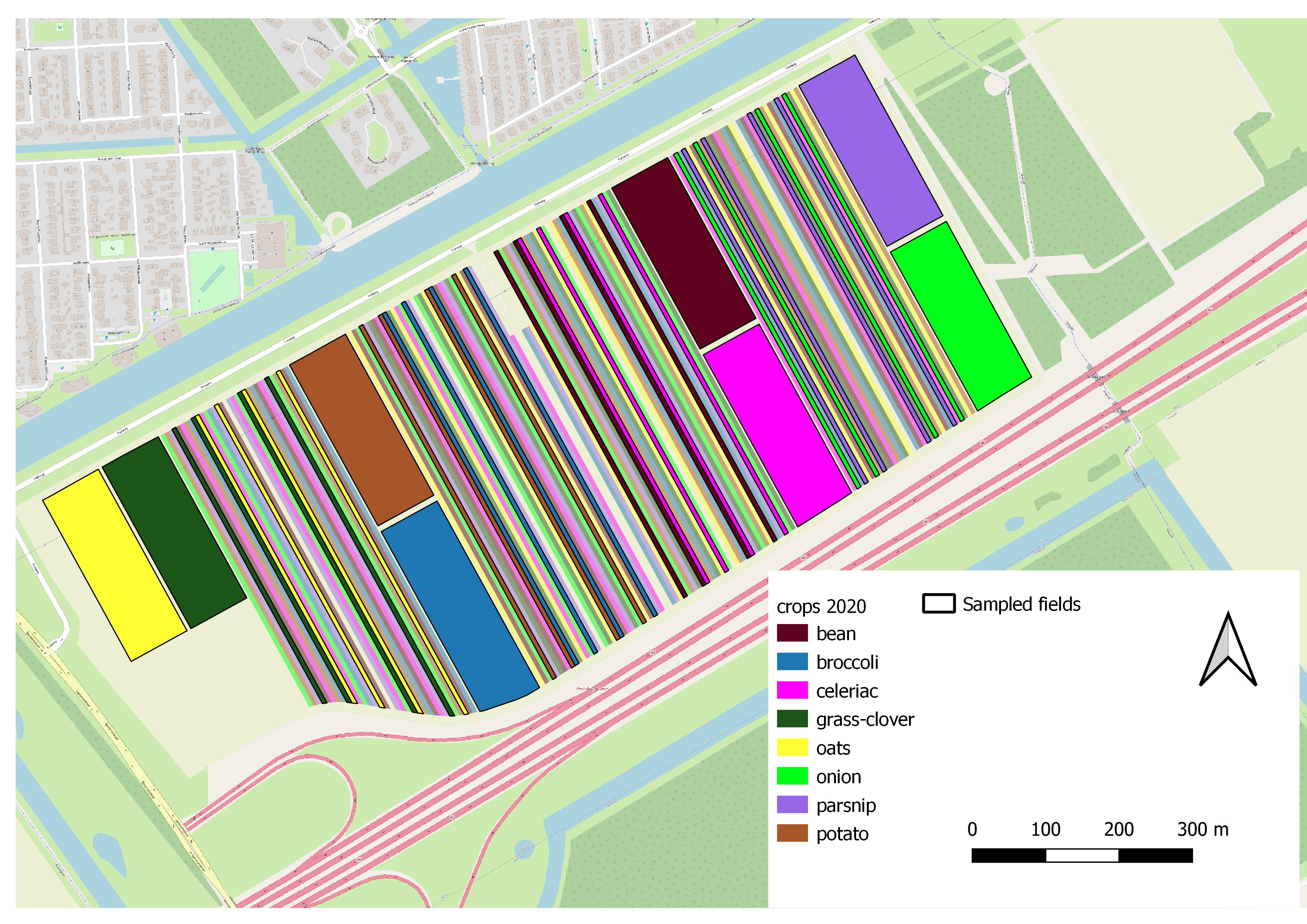

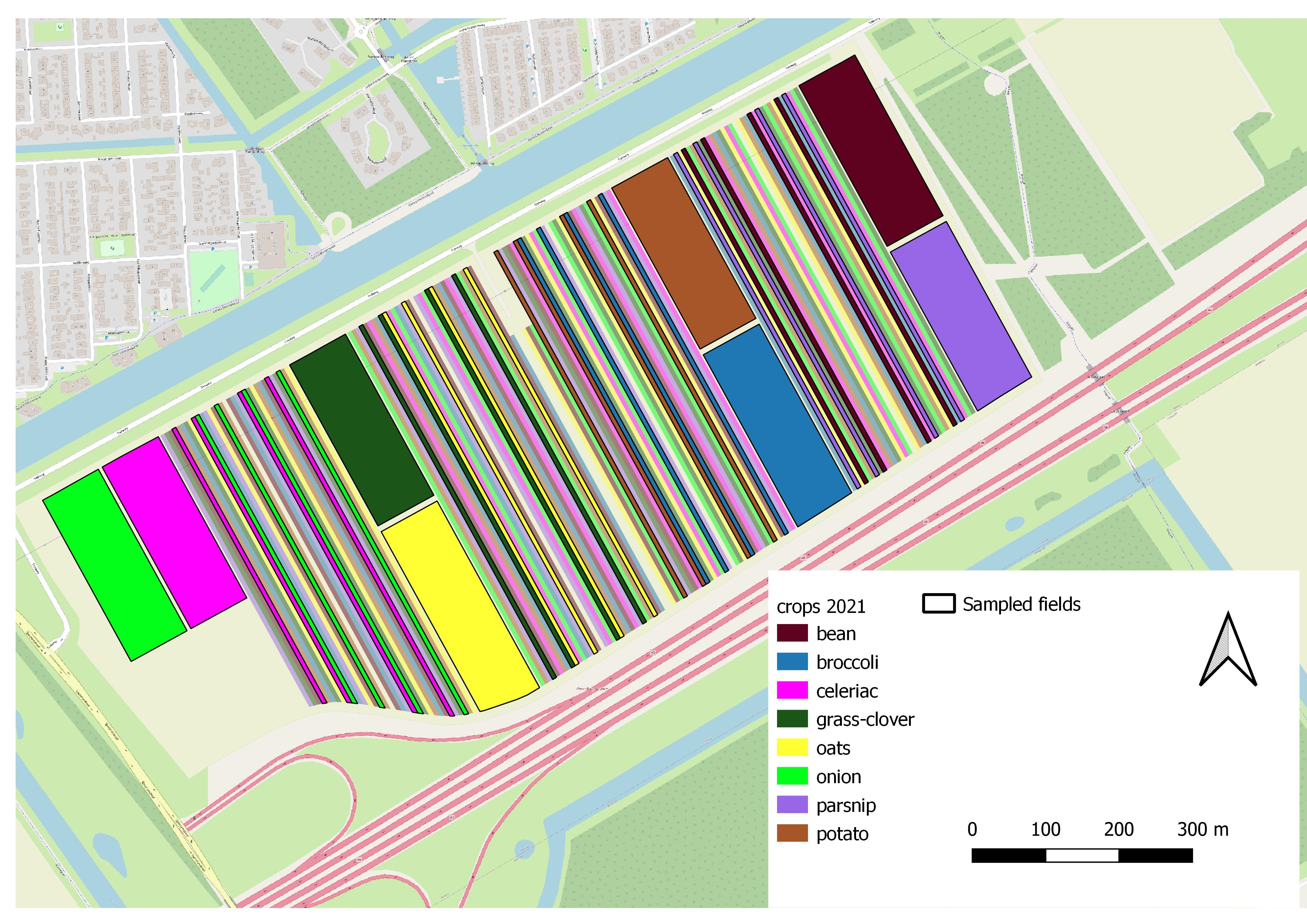

**A Almere (continued)**

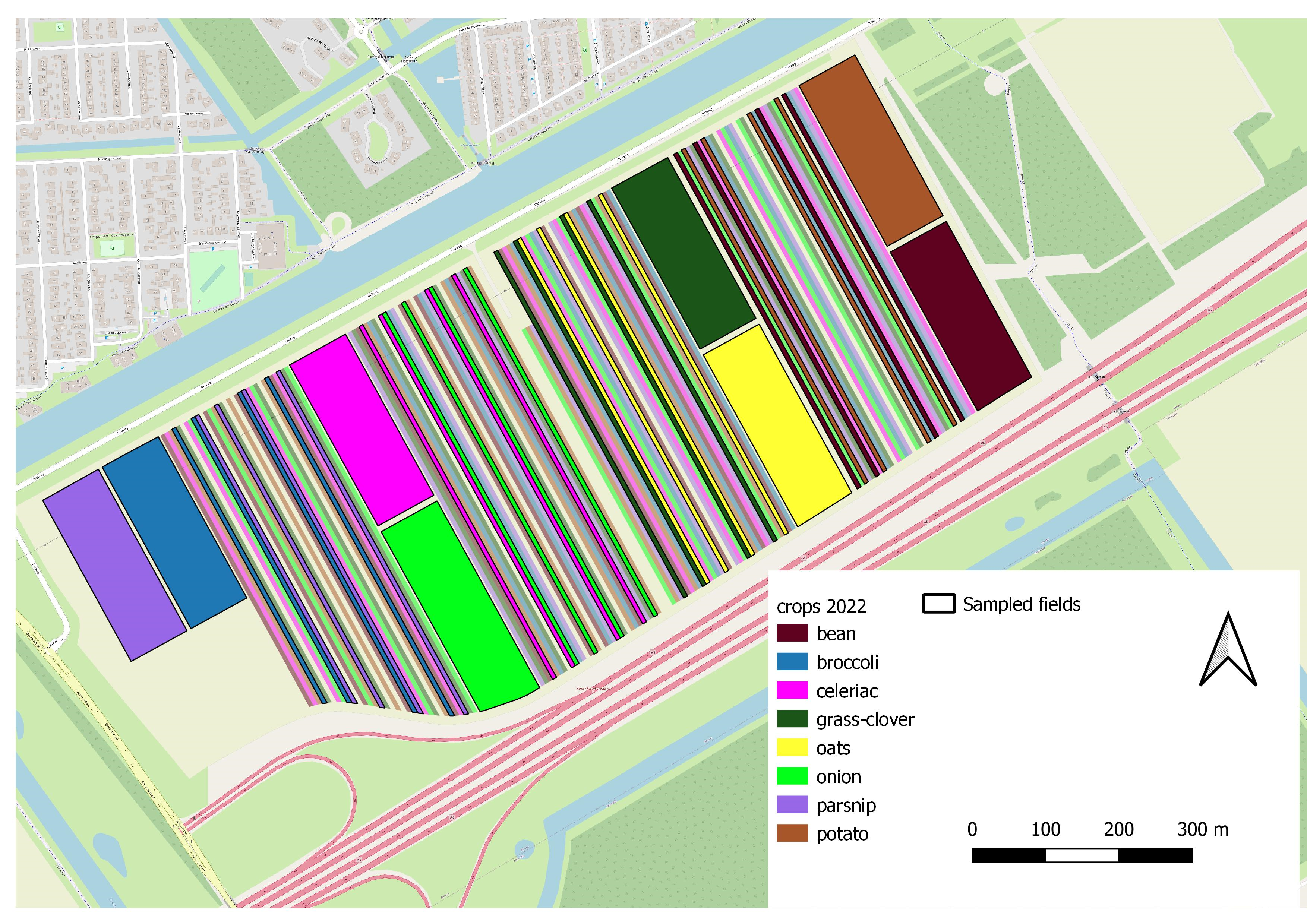

**B Lelystad**

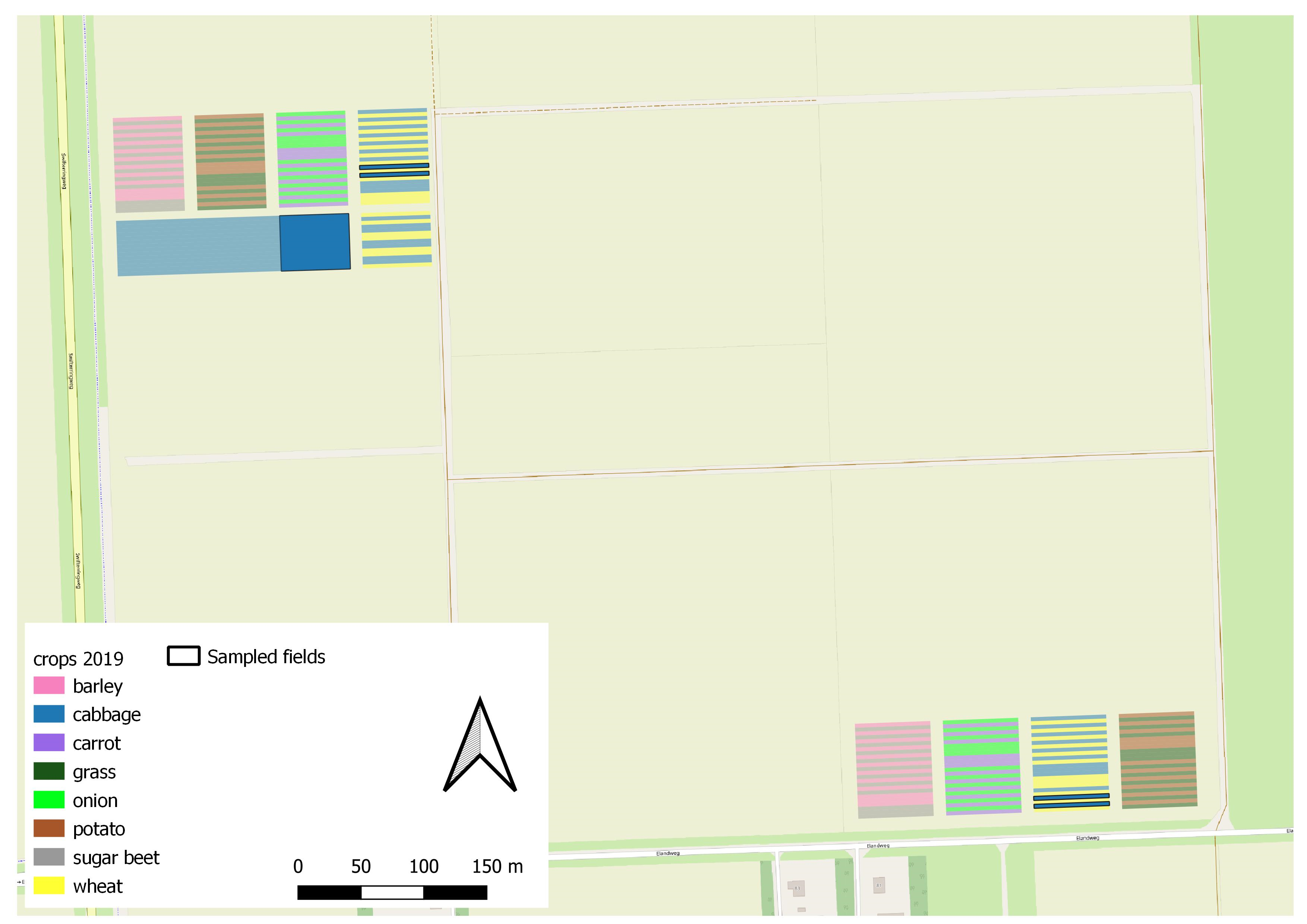

**B Lelystad (continued)**

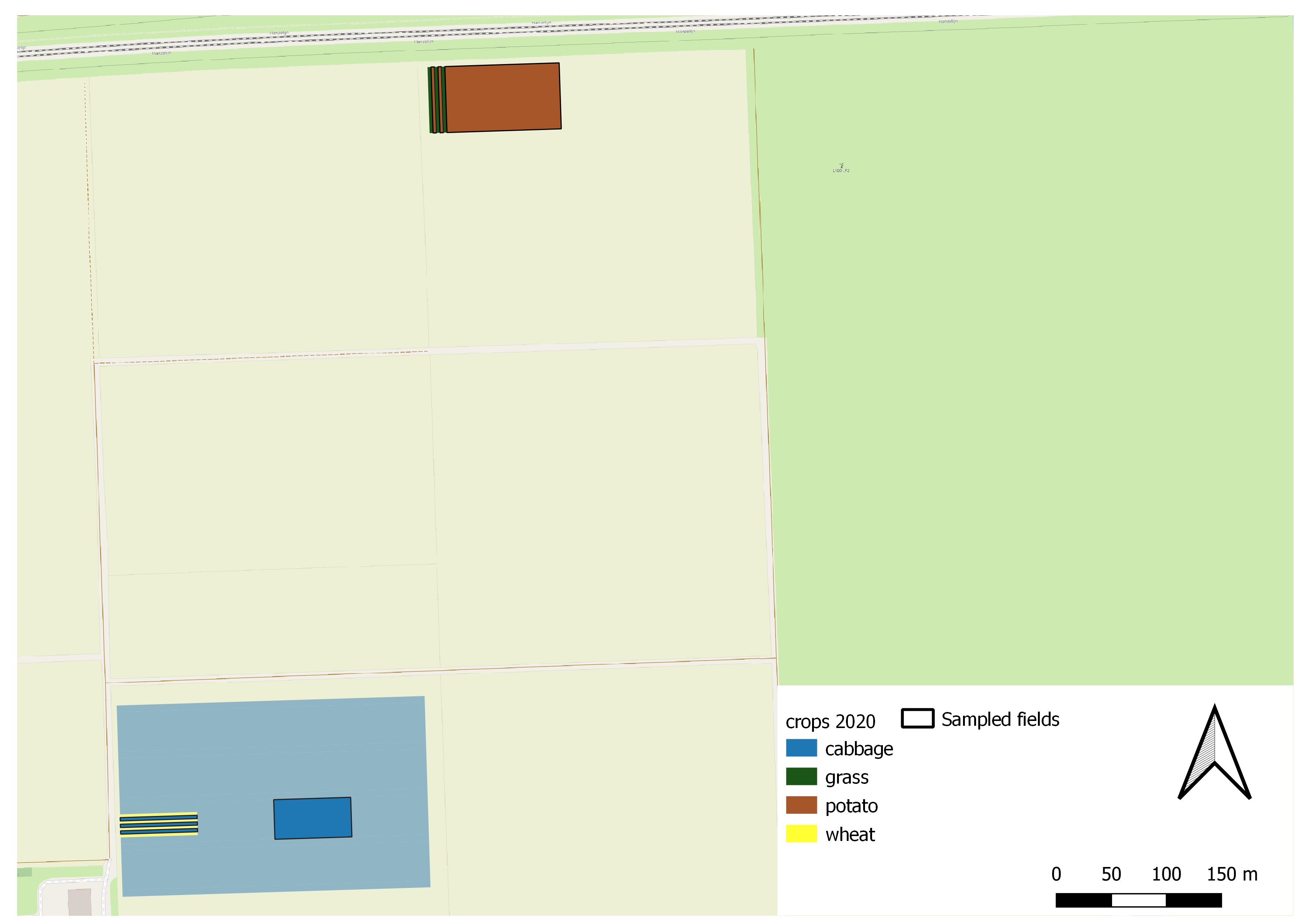

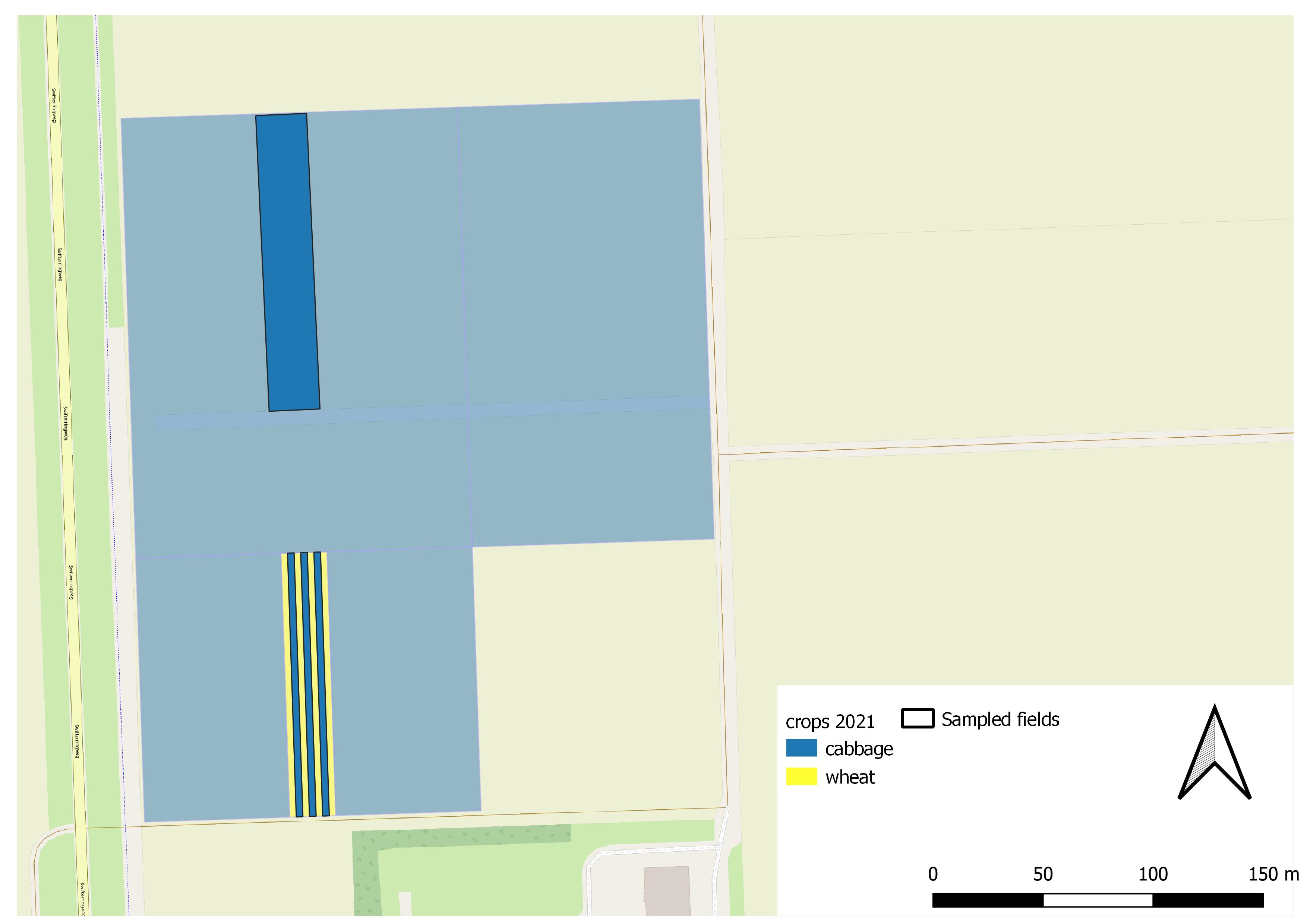

**C Valthermond**

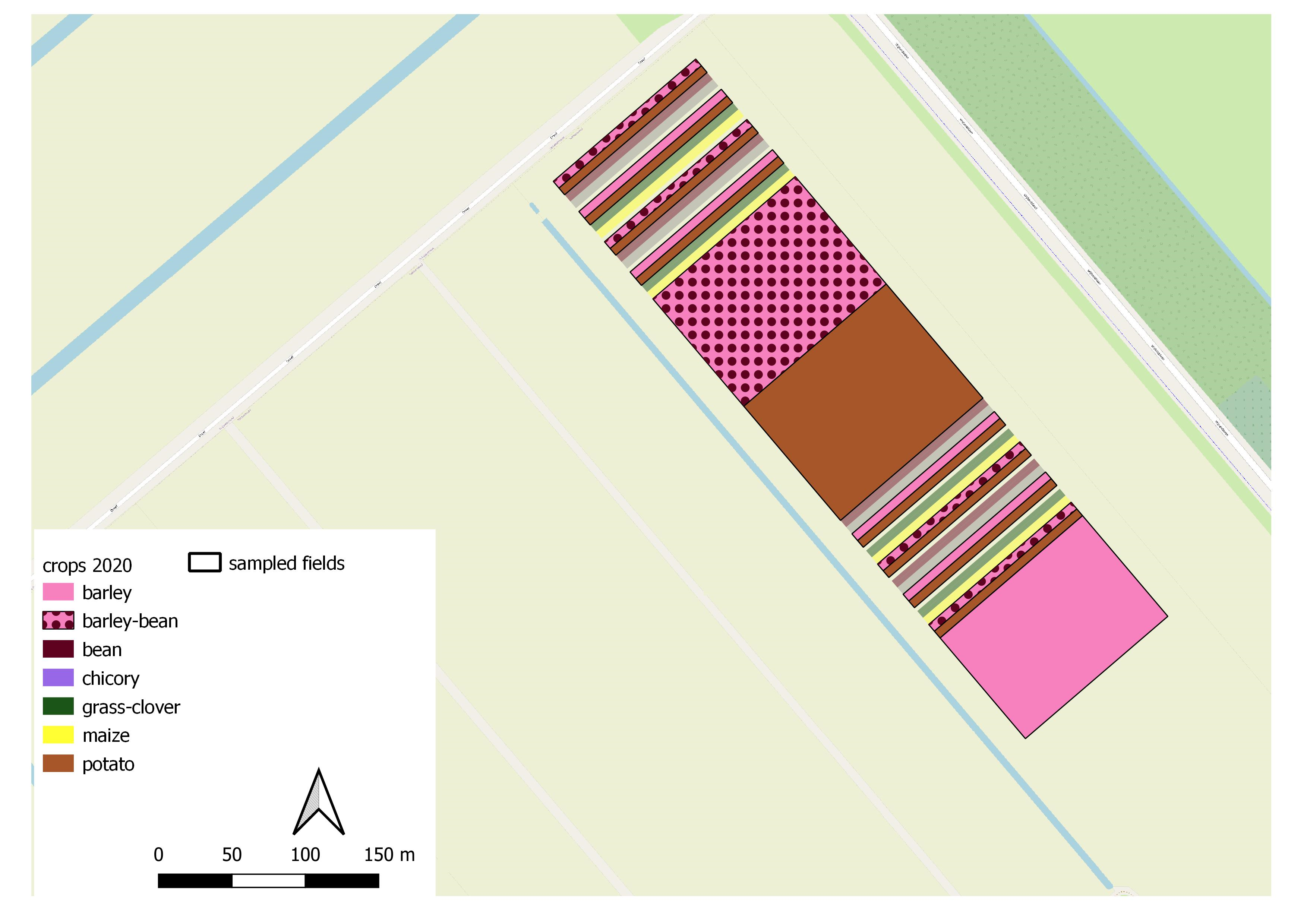

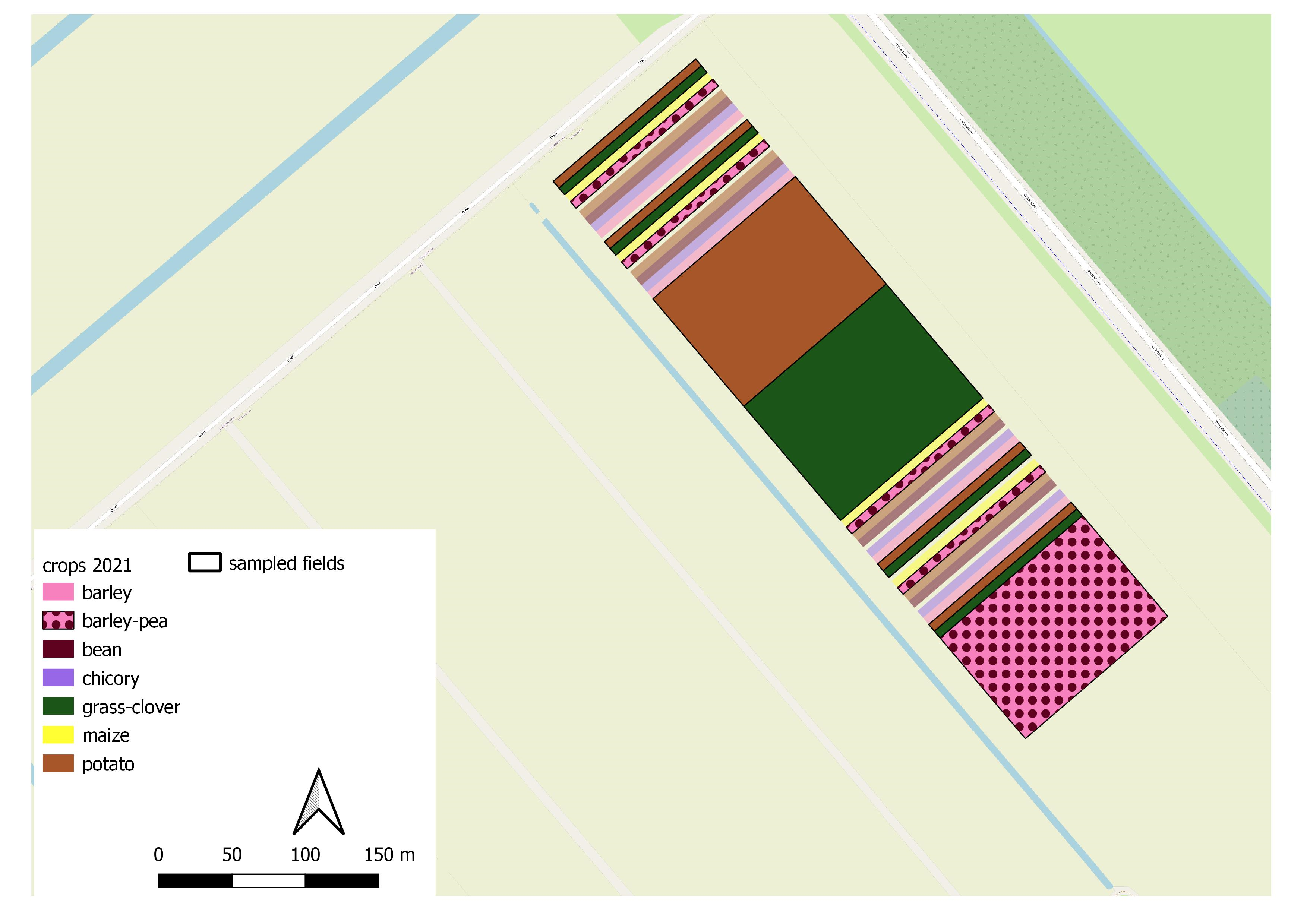

**D Wageningen**

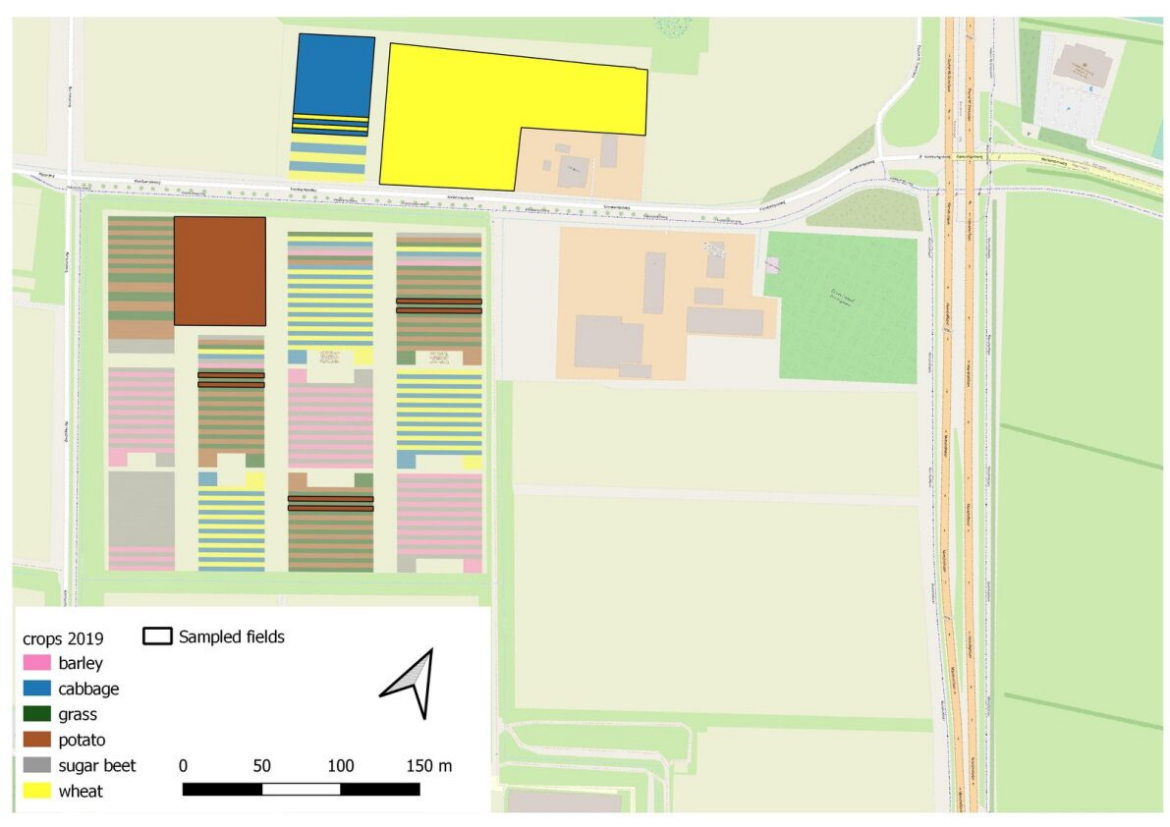

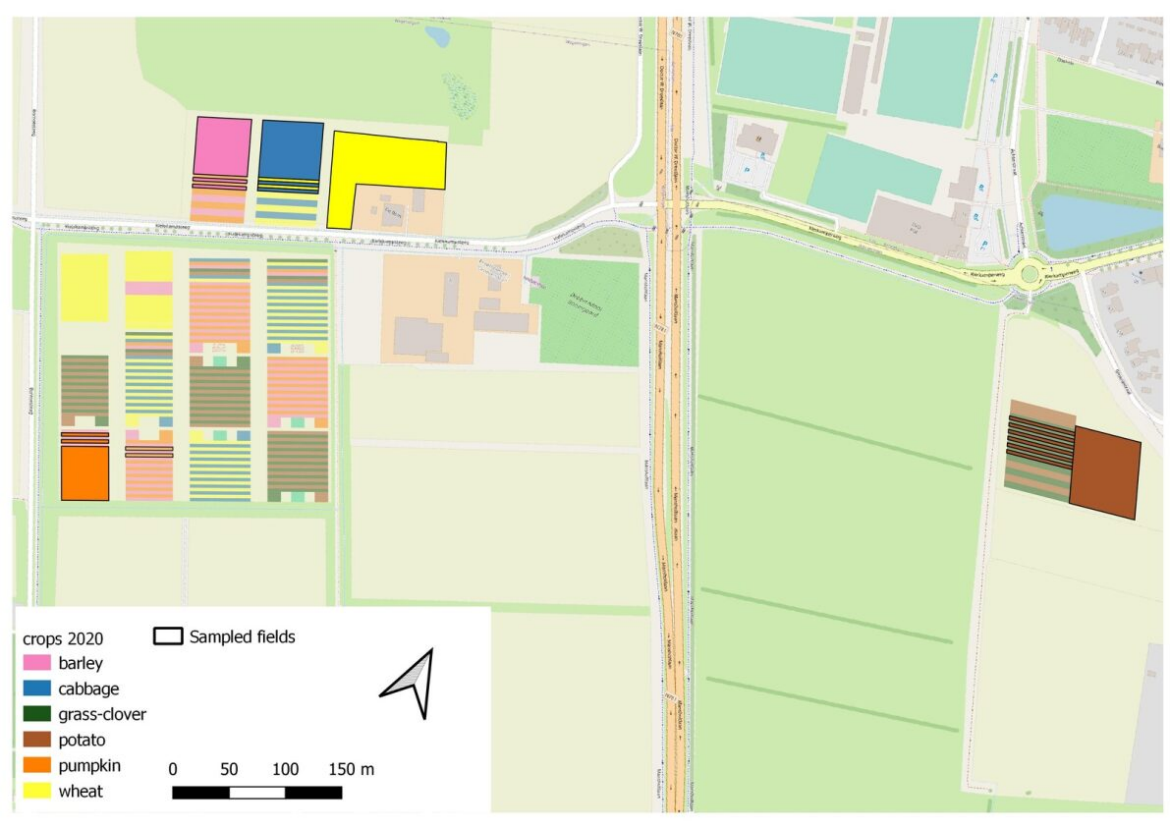

**D Wageningen (continued)**

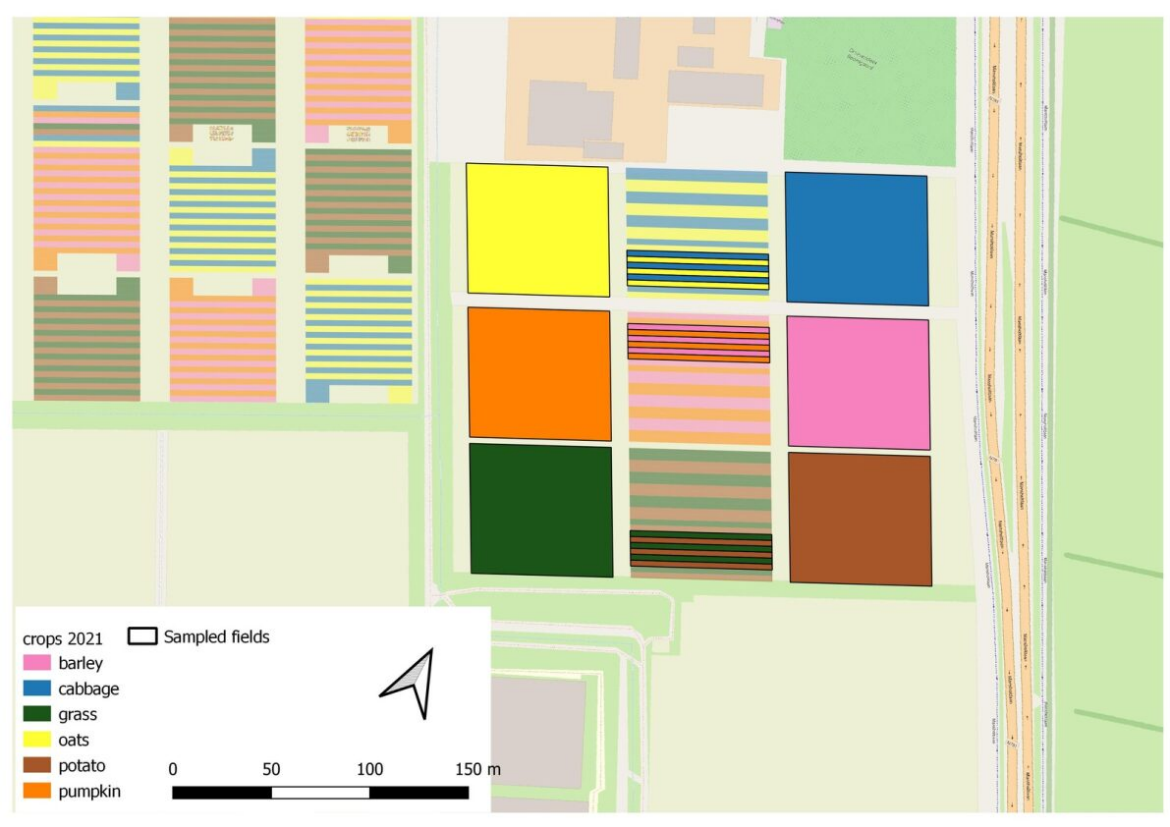

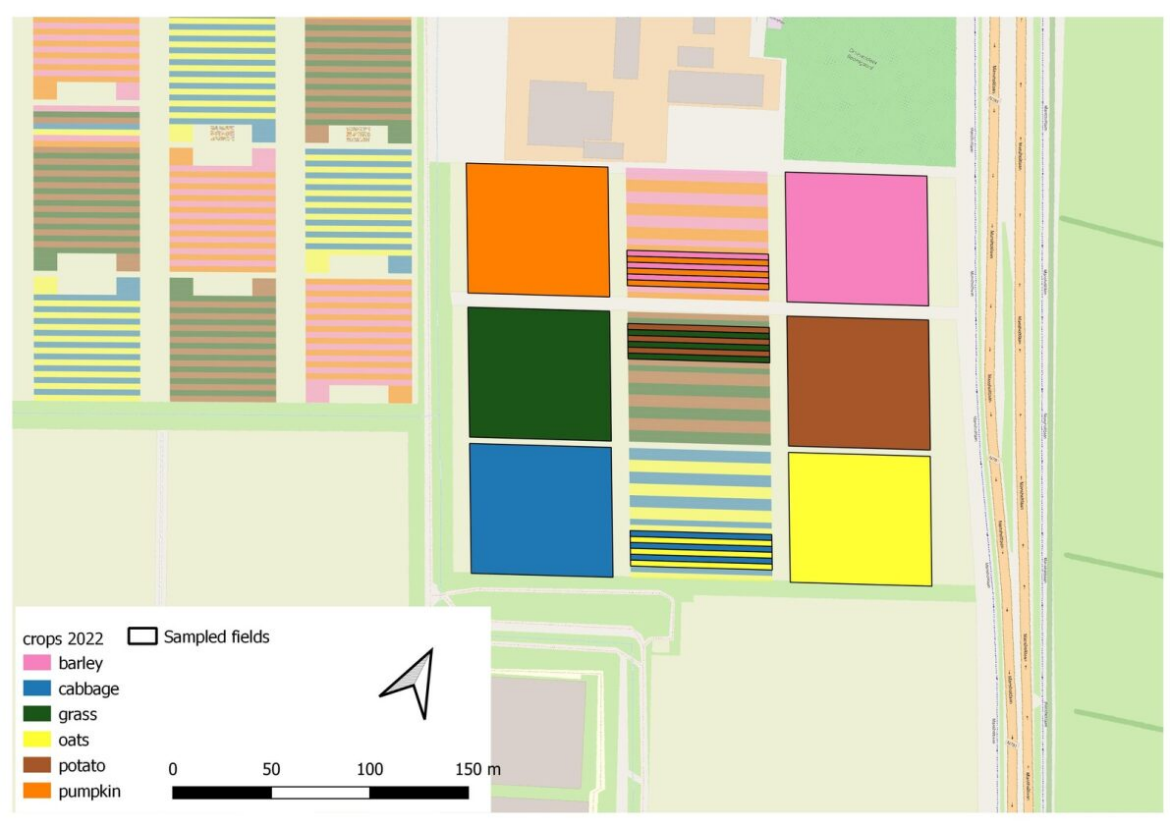

**Fig. S10. Field maps of experimental lay-out per year and location.** Field maps are shown in chronological order (oldest to newest) per location: Almere **(a)**, Lelystad **(b)**, Valthermond **(c)** and Wageningen **(d)**. Colors indicate distinct crops, whereas diagonal lines indicate a crop mixture (only for Valthermond). Brighter colours with black lining around the field indicate areas in which samples included in this study were taken.

**Supplementary tables**

**Table S1.** All ground beetle species found among the four locations, and their species codes as used in figures S3-S5 and S9.

| **Subfamily** | **Genus** | **Species** | **Species code** |
| --- | --- | --- | --- |
| Carabinae | Carabus | *Carabus granulatus* | CaraGran |
|  | Nebria | *Nebria brevicollis* | NebrBrev |
|  |  | *Nebria salina* | NebrSali |
|  | Notiophilus | *Notiophilus aquaticus* | NotiAqua |
|  |  | *Notiophilus biguttatus* | NotiBigu |
|  |  | *Notiophilus palustris* | NotiPalu |
| Harpalinae | Acupalpus | *Acupalpus meridianus* | AcupMeri |
|  | Agonum | *Agonum muelleri* | AgonMuel |
|  | Amara | *Amara aenea* | AmarAene |
|  |  | *Amara anthobia* | AmarAnth |
|  |  | *Amara apricaria* | AmarApri |
|  |  | *Amara aulica* | AmarAuli |
|  |  | *Amara bifrons* | AmarBifr |
|  |  | *Amara communis* | AmarComm |
|  |  | *Amara consularis* | AmarCons |
|  |  | *Amara famelica* | AmarFame |
|  |  | *Amara familiaris* | AmarFami |
|  |  | *Amara fulva* | AmarFulv |
|  |  | *Amara ovata* | AmarOvat |
|  |  | *Amara plebeja* | AmarPleb |
|  |  | *Amara similata* | AmarSimi |
|  |  | *Amara spreta* | AmarSpre |
|  |  | *Amara tibialis* | AmarTibi |
|  | Anchomenus | *Anchomenus dorsalis* | AnchDors |
|  | Anisodactylus | *Anisodactylus binotatus* | AnisBino |
|  | Badister | *Badister bullatus* | BadiBull |
|  |  | *Badister sodalist* | BadiSoda |
|  | Bradycellus | *Bradycellus harpalinus* | BradHarp |
|  | Calathus | *Calathus cinctus* | CalaCinc |
|  |  | *Calathus erratus* | CalaErra |
|  |  | *Calathus fuscipes* | CalaFusc |
|  |  | *Calathus melanocephalus* | CalaMela |
|  |  | *Calathus rotundicollis* | CalaRotu |
|  | Harpalus | *Harpalus affinis* | HarpAffi |
|  |  | *Harpalus distinguendus* | HarpDist |
|  |  | *Harpalus griseus* | HarpGris |
|  |  | *Harpalus rubripes* | HarpRubr |
|  |  | *Harpalus rufipes* | HarpRufi |
|  |  | *Harpalus signaticornis* | HarpSign |
|  |  | *Harpalus tardus* | HarpTard |
|  | Microlestes | *Microlestes minutulus* | MicrMinu |
|  | Oxypselaphus | *Oxypselaphus obscurus* | OxypObsc |
|  | Panagaeus | *Panagaeus bipustulatus* | PanaBipu |
|  | Poecilus | *Poecilus cupreus* | PoecCupr |
|  |  | *Poecilus versicolor* | PoecVers |
|  | Pterostichus | *Pterostichus anthracinus* | PterAnth |
|  |  | *Pterostichus melanarius* | PterMela |
|  |  | *Pterostichus niger* | PterNige |
|  |  | *Pterostichus strenuus* | PterStre |
|  |  | *Pterostichus vernalis* | PterVern |
|  | Stenolophus | *Stenolophus teutonus* | StenTeut |
|  | Stomis | *Stomis pumicatus* | StomPumi |
|  | Syntomus | *Syntomus truncatellus* | SyntTrun |
| Loricerinae | Loricera | *Loricera pilicornis* | LoriPili |
| Scaritinae | Broscus | *Broscus cephalotes* | BrosCeph |
|  | Clivina | *Clivina collaris* | ClivColl |
|  |  | *Clivina fossor* | ClivFoss |
|  | Dyschirius | *Dyschirius globosus* | DyscGlob |
| Trechinae | Asaphidion | *Asaphidion flavipes* | AsapFlav |
|  | Bembidion | *Bembidion aenea* | BembAene |
|  |  | *Bembidion biguttatum* | BembBigu |
|  |  | *Bembidion femoratum* | BembFemo |
|  |  | *Bembidion lampros* | BembLamp |
|  |  | *Bembidion lunulatum* | BembLunu |
|  |  | *Bembidion obtusum* | BembObtu |
|  |  | *Bembidion proprans* | BembProp |
|  |  | *Bembidion quadrimaculatum* | BembQuadrim |
|  |  | *Bembidion tetracolum* | BembTetr |
|  | Blemus | *Blemus discus* | BlemDisc |
|  | Trechoblemus | *Trechoblemus micros* | TrecMicr |
|  | Trechus | *Trechus obtusus* | TrecObtu |
|  |  | *Trechus quadristriatus* | TrecQuad |

**Table S2.** Total number of ground beetles caught per species (or genus), per location. The number of yearseries is given per location in brackets. For some locations, ground beetles were identified up to genus level, these are underlined. “N/A” indicates that this taxa was identified to a different taxonomic level for the specific location. Locations are indicated with abbreviations (Al=Almere; Le = Lelystad; Va = Valthermond; Wa = Wageningen). Rarity indicates the rarity of the species according to waarneming.nl (1 = common, 2 = relatively common, 3 = rare, 4 = very rare). Affinity indicates the habitat affinity group as by table A.1 of Turin et al. (2022), a * here indicates that species are more eurytopic.

|  | **Species** | **Rarity** | **Affinity** | **Al**  **(183)** | **Le**  **(27)** | **Va**  **(72)** | **Wa**  **(179)** | **Total** |
| --- | --- | --- | --- | --- | --- | --- | --- | --- |
|  | **All Carabidae** |  |  | **40153** | **3777** | **1126** | **3052** | **48108** |
|  | **Total Amara** |  |  | **171** | **0** | **27** | **392** | **590** |
| 1 | *Amara aenea* | 1 | Grassland* | 15 | 0 | 8 | 163 | 186 |
| 2 | *Amara anthobia* | 2 | Forest* | 0 | 0 | 3 | 0 | 3 |
| 3 | *Amara apricaria* | 1 | Ruderal* | 0 | 0 | 0 | 19 | 19 |
| 4 | *Amara aulica* | 2 | Grassland | 1 | 0 | 0 | 1 | 2 |
| 5 | *Amara bifrons* | 2 | Ruderal* | 1 | 0 | 1 | 36 | 38 |
| 6 | *Amara communis* | 1 | Grassland* | 0 | 0 | 0 | 1 | 1 |
| 7 | *Amara consularis* | 2 | Ruderal | 0 | 0 | 0 | 54 | 54 |
| 8 | *Amara familiaris* | 1 | Grassland* | 124 | 0 | 0 | 5 | 129 |
| 9 | *Amara fulva* | 2 | Ruderal | 0 | 0 | 6 | 97 | 103 |
| 10 | *Amara ovata* | 2 | Grassland | 0 | 0 | 9 | 0 | 9 |
| 11 | *Amara plebeja* | 1 | Heathland* | 1 | 0 | 0 | 0 | 1 |
| 12 | *Amara similata* | 1 | Ruderal* | 24 | 0 | 0 | 4 | 28 |
| 13 | *Amara spreta* | 1 | Dunes* | 0 | 0 | 0 | 11 | 11 |
| 14 | *Amara tibialis* | 2 | Grassland* | 0 | 0 | 0 | 1 | 1 |
|  | *Amara sp.* |  |  | 5 | N/A | N/A | N/A | 5 |
|  | **Total Anchomenus** |  |  | **151** | **1** | **161** | **8** | **321** |
| 15 | *Anchomenus dorsalis* | 1 | Ruderal | 151 | 1 | 161 | 8 | 321 |
|  | **Total Bembidion** |  |  | **753** | **407** | **32** | **150** | **1342** |
| 16 | *Bembidion aeneum* | 2 | Wetland* | 0 | 1 | N/A | 0 | 1 |
| 17 | *Bembidion biguttatum* | 1 | Ruderal | 7 | 0 | N/A | 0 | 7 |
| 18 | *Bembidion femoratum* | 1 | Wetland | 0 | 0 | N/A | 30 | 30 |
| 19 | *Bembidion lampros* | 1 | Heathland* | 3 | 1 | N/A | 21 | 25 |
| 20 | *Bembidion lunulatum* | 1 | Ruderal* | 3 | 0 | N/A | 0 | 3 |
| 21 | *Bembidion obtusum* | 2 | Ruderal* | 4 | 0 | N/A | 0 | 4 |
| 22 | *Bembidion proprans* | 1 | Ruderal | 4 | 0 | N/A | 23 | 27 |
| 23 | *Bembidion quadrim.* | 1 | Ruderal | 87 | 8 | N/A | 33 | 128 |
| 24 | *Bembidion tetracolum* | 1 | Ruderal* | 345 | 397 | N/A | 43 | 785 |
|  | *Bembidion sp.* |  |  | 300 | N/A | 32 | N/A | 332 |
|  | **Total Blemus** |  |  | **28** | **45** | **0** | **0** | **79** |
| 25 | *Blemus discus* | 2 | Ruderal | 28 | 45 | 0 | 0 | 79 |
|  | **Total Calathus** |  |  | **0** | **0** | **95** | **203** | **298** |
| 26 | *Calathus cinctus* | 2 | Ruderal* | 0 | 0 | 20 | 73 | 93 |
| 27 | *Calathus erratus* | 2 | Heathland* | 0 | 0 | 6 | 87 | 93 |
| 28 | *Calathus fuscipes* | 1 | Grassland* | 0 | 0 | 2 | 0 | 2 |
| 29 | *Calathus melanocephalus* | 1 | Heathland* | 0 | 0 | 67 | 41 | 108 |
| 30 | *Calathus rotundicollis* | 2 | Forest | 0 | 0 | 0 | 2 | 2 |
|  | **Total Clivina** |  |  | **54** | **11** | **4** | **62** | **131** |
| 31 | *Clivina collaris* | 1 | Ruderal* | 0 | 2 | 2 | 28 | 32 |
| 32 | *Clivina fossor* | 1 | Ruderal* | 49 | 9 | 2 | 34 | 94 |
|  | *Clivina sp.* |  |  | 5 | N/A | N/A | N/A | 5 |
|  | **Total Harpalus** |  |  | **2496** | **19** | **481** | **1161** | **4157** |
| 33 | *Harpalus affinis* | 1 | Ruderal* | 131 | 3 | 1 | 30 | 165 |
| 34 | *Harpalus distinguendus* | 2 | Ruderal | 0 | 0 | N/A | 2 | 2 |
| 35 | *Harpalus griseus* | 2 | Forest | 9 | 0 | 2 | 4 | 15 |
| 36 | *Harpalus rubripes* | 2 | Grassland | 0 | 1 | N/A | 1 | 2 |
| 37 | *Harpalus rufipes* | 1 | Ruderal* | 1409 | 15 | 168 | 1104 | 2696 |
| 38 | *Harpalus signaticornis* | 4 | Grassland | 0 | 0 | N/A | 5 | 5 |
| 39 | *Harpalus tardus* | 1 | Grassland* | 0 | 0 | 2 | 15 | 17 |
|  | *Harpalus sp.* |  |  | 947 | N/A | 308 | N/A | 1255 |
|  | **Total Loricera** |  |  | **22** | **6** | **20** | **21** | **69** |
| 40 | *Loricera pilicornis* | 1 | Ruderal* | 22 | 6 | 20 | 21 | 69 |
|  | **Total Nebria** |  |  | **362** | **2** | **0** | **33** | **397** |
| 41 | *Nebria brevicollis* | 1 | Forest* | 320 | 2 | 0 | 16 | 338 |
| 42 | *Nebria salina* | 2 | Heathland | 0 | 0 | 0 | 17 | 17 |
|  | *Nebria sp.* |  |  | 42 | N/A | N/A | N/A | 42 |
|  | **Total Poecilus** |  |  | **4649** | **235** | **82** | **49** | **5015** |
| 43 | *Poecilus cupreus* | 1 | Ruderal | 2876 | 233 | N/A | 24 | 3133 |
| 44 | *Poecilus versicolor* | 1 | Heathland | 4 | 2 | N/A | 25 | 32 |
|  | *Poecilus sp.* |  |  | 1769 | N/A | 82 | N/A | 1851 |
|  | **Total Pterostichus** |  |  | **31041** | **2968** | **217** | **769** | **34995** |
| 45 | *Pterostichus anthracinus* | 2 | Forest | 1 | 0 | N/A | 0 | 1 |
| 46 | *Pterostichus melanarius* | 1 | Ruderal | 5556 | 2731 | 144 | 763 | 9193 |
| 47 | *Pterostichus niger* | 1 | Heathland* | 167 | 228 | 5 | 1 | 401 |
| 48 | *Pterostichus strenuus* | 1 | Forest* | 2 | 0 | N/A | 0 | 2 |
| 49 | *Pterostichus vernalis* | 1 | Ruderal* | 57 | 9 | N/A | 5 | 73 |
|  | *Pterostichus sp.* |  |  | 25258 | N/A | 68 | N/A | 25326 |
|  | **Total Trechus** |  |  | **374** | **76** | **0** | **143** | **593** |
| 50 | *Trechus obtusus* | 1 | Grassland* | 2 | 0 | 0 | 18 | 20 |
| 51 | *Trechus quadristriatus* | 1 | Ruderal* | 69 | 76 | 0 | 125 | 270 |
|  | *Trechus sp.* |  |  | 303 | N/A | N/A | N/A | 303 |
|  | **Total other Carabids** |  |  | **52** | **7** | **7** | **61** | **127** |
| 52 | *Acupalpus meridianus* | 2 | Ruderal | 8 | 3 | 0 | 1 | 12 |
|  | *Acupalpus sp.* |  |  | 2 | N/A | N/A | N/A | 2 |
| 53 | *Agonum muelleri* | 1 | Ruderal | 5 | 3 | 7 | 5 | 20 |
| 54 | *Anisodactylus binotatus* | 1 | Ruderal* | 6 | 0 | 0 | 1 | 7 |
| 55 | *Badister bullatus* | 1 | Dunes | 2 | 0 | 0 | 1 | 3 |
| 56 | *Badister sodalis* | 2 | Heathland | 3 | 0 | 0 | 0 | 3 |
| 57 | *Bradycellus harpalinus* | 1 | Heathland | 1 | 0 | 0 | 0 | 1 |
| 58 | *Broscus cephalotes* | 1 | Dunes | 0 | 0 | 0 | 35 | 35 |
| 59 | *Carabus granulatus* | 1 | Heathland | 2 | 0 | 0 | 0 | 2 |
|  | *Carabus sp.* |  |  | 1 | N/A | N/A | N/A | 1 |
| 60 | *Dyschirius globosus* | 1 | Heathland | 0 | 0 | 0 | 10 | 10 |
| 61 | *Microlestes minutulus* | 3 | Dunes* | 0 | 0 | 0 | 1 | 1 |
| 62 | *Notiophilus aquaticus* | 1 | Heathland | 0 | 0 | 0 | 1 | 1 |
| 63 | *Notiophilus palustris* | 1 | Grassland* | 2 | 0 | 0 | 0 | 2 |
| 64 | *Oxypselaphus obscurus* | 1 | Heathland | 3 | 0 | 0 | 0 | 3 |
| 65 | *Stenolophus teutonus* | 1 | Ruderal* | 0 | 0 | 0 | 2 | 2 |
| 66 | *Stomis pumicatus* | 1 | Forest | 3 | 0 | 0 | 0 | 3 |
| 67 | *Syntomus foveatus* | 1 | Dunes* | 0 | 0 | 0 | 1 | 1 |
| 68 | *Trechoblemus micros* | 2 | Ruderal | 4 | 0 | 0 | 0 | 4 |
|  | Unknown carabidae |  |  | 10 | 1 | 0 | 3 | 14 |

**Table S3.** Abundances of the twelve most abundant ground beetle genera in monoculture and strip cropping fields in four locations. The first value indicates the estimated mean, in brackets the confidence interval and letters indicate significant difference between monocultures and strips per location. A strip indicates that there was no significant difference for the genus for that location. Dark cells indicate that two or less individuals were found at this location, and the location was excluded from the model. At Wageningen, we only found few *Anchomenus* in the monoculture, whereas we found none in the strip cropped field. As there was no variation in the strip cropped field, this location could not be included in the model for this genus and the mean here indicates the actual mean. The model for Pterostichus did not fit well when the data from 2020 Almere were included, as catches were much higher in this year than in the other years. Therefore, we conducted separate analyses for2020 and 2021-2022.

|  | **Almere** | | **Lelystad** | | **Valthermond** | | **Wageningen** | |
| --- | --- | --- | --- | --- | --- | --- | --- | --- |
| **Genus** | **Mono** | **Strip** | **Mono** | **Strip** | **Mono** | **Strip** | **Mono** | **Strip** |
| *Amara* | 0.44  (0.17-1.15)  B | 0.25  (0.09-0.67)  A |  |  | 0.11  (0.02-0.57)  - | 0.13  (0.03-0.55)  - | 0.51  (0.20-1.33)  - | 0.76  (0.29-2.00)  - |
| *Anchomenus* | 0.06  (0.02-0.20)  A | 0.20  (0.06-0.64)  B |  |  | 0.91  (0.18-4.70)  - | 1.89  (0.41-8.78)  - | 0.04  N/A | 0.00  N/A |
| *Bembidion* | 2.91  (2.07-4.10)  A | 4.80  (3.44-6.70)  B | 6.32  (3.30-12.1)  - | 4.35  (2.12-8.93)  - | 0.46  (0.20-1.06)  - | 0.52  (0.28-0.98)  - | 0.53  (0.35-0.81)  - | 0.56  (0.34-0.91)  - |
| *Blemus* | 0.15  (0.04-0.65)  - | 0.12  (0.03-0.51)  - | 0.73  (0.16-3.36)  - | 0.41  (0.08-2.03)  - |  |  |  |  |
| *Calathus* |  |  |  |  | 2.24  (1.03-4.89)  B | 1.22  (0.57-2.60)  A | 0.50  (0.27-0.93)  - | 0.44  (0.22-0.85)  - |
| *Clivina* | 0.18  (0.08-0.40)  - | 0.15  (0.06-0.34)  - | 0.18  (0.05-0.62)  - | 0.05  (0.01-0.33)  - | 0.10  (0.03-0.42)  - | 0.02  (0.00-0.14)  - | 0.18  (0.08-0.39)  - | 0.11  (0.04-0.28)  - |
| *Harpalus* | 11.2  (6.50-19.3)  A | 15.1  (8.75-26.0)  B | 0.52  (0.21-1.32)  - | 0.62  (0.23-1.68  - | 5.04  (2.51-10.1)  - | 4.40  (2.30-8.41)  - | 4.43  (2.58-7.60)  A | 6.12  (3.50-10.7)  B |
| *Loricera* | 0.05  (0.01-0.32)  - | 0.02  (0.00-0.17)  - | 0.27  (0.03-2.89)  - | 0.14  (0.01-1.92)  - | 0.24  (0.03-1.79)  - | 0.08  (0.01-0.60)  - | 0.03  (0.00-0.19)  - | 0.06  (0.01-0.41)  - |
| *Nebria* | 0.70  (0.23-2.17)  A | 1.30  (0.42-4.00)  B |  |  |  |  | 0.10  (0.03-0.31)  - | 0.06  (0.02-0.24)  - |
| *Poecilus* | 19.1  (13.8-26.5)  - | 21.8  (15.7-30.3)  - | 1.83  (0.98-3.39)  - | 1.54  (0.76-3.11)  - | 1.03  (0.52-2.04)  - | 0.87  (0.47-1.61)  - | 0.17  (0.10-0.29)  - | 0.18  (0.10-0.34)  - |
| *2020*  *Pterostichus*  *2021-22* | 434  (325-581)  B  47.2  (31.2-71.5)  - | 307  (230-411)  A  64.1  (41.6-98.7)  - | 35.1  (16.9-72.6)  - | 30.0  (13.3-68.0)  - | 3.93  (2.09-7.41)  B | 1.70  (1.00-2.88)  A | `3.02  (2.01-4.55)  - | 2.49  (1.58-3.91)  - |
| *Trechus* | 1.25  (0.69-2.26)  B | 0.73  (0.40-1.33)  A | 3.54  (1.37-9.12)  B | 0.91  (0.30-2.79)  A |  |  | 0.55  (0.31-0.98)  - | 0.32  (0.16-0.64)  - |

**Table S4. Effect of crop configuration on ground beetle community composition.** Results from permanova analyses using Hellinger’s transformation for data from the three locations with species level data. “Crop species” is a nested variable within years, as these differed among years. Years were nested in locations, as the years that were studied differed among locations. P-values in bold typeset indicate significant effects (α = 0.05).

| Location | Predictor | Df | Sum  Sq | R2 | F | P |
| --- | --- | --- | --- | --- | --- | --- |
| Almere | Crop configuration | 1 | 0.20 | 0.01 | 2.34 | **0.025** |
|  | Year | 1 | 2.48 | 0.13 | 28.6 | **0.001** |
|  | Year : Crop species | 14 | 5.66 | 0.30 | 4.68 | **0.001** |
|  | Crop configuration : Year | 1 | 0.13 | 0.01 | 1.50 | 0.138 |
|  | Crop configuration : Year : Crop species | 14 | 2.88 | 0.15 | 2.38 | **0.001** |
|  | *Residual* | 88 | *7.61* | *0.40* |  |  |
|  | *Total* | 119 | *19.0* | *1.00* |  |  |
| Lelystad | Crop configuration | 1 | 0.03 | 0.01 | 0.68 | 0.534 |
|  | Year | 2 | 1.78 | 0.55 | 18.9 | **0.001** |
|  | Year : Crop species | 1 | 0.12 | 0.04 | 2.45 | 0.082 |
|  | Crop configuration : Year | 2 | 0.26 | 0.08 | 2.80 | **0.026** |
|  | Crop configuration : Year : Crop species | 1 | 0.14 | 0.04 | 2.91 | **0.039** |
|  | *Residual* | *19* | *0.90* | *0.28* |  |  |
|  | *Total* | *26* | *3.22* | *1.00* |  |  |
| Wageningen | Crop configuration | 1 | 1.48 | 0.02 | 4.30 | **0.001** |
|  | Year | 3 | 6.40 | 0.07 | 6.22 | **0.001** |
|  | Year : Crop species | 16 | 27.7 | 0.29 | 5.05 | **0.001** |
|  | Crop configuration : Year | 3 | 1.90 | 0.02 | 1.85 | **0.006** |
|  | Crop configuration : Year : Crop species | 16 | 11.8 | 0.12 | 2.15 | **0.001** |
|  | *Residual* | *138* | *47.3* | *0.49* |  |  |
|  | *Total* | *177* | *96.6* | *1.00* |  |  |

**Table S5. Effect of crop configuration and crop species on ground beetle community composition.** Results from pairwise permanova analyses for crop pairs pumkin-barley, cabbage-oat, and potato-grass in 2021 (a) and 2022 (b) in Wageningen. Values show F-values for the comparison between the crop configurations and crops in crossing rows and columns. Bold numbers indicate significant differences between combinations of crop configurations and crops (α = 0.05).

**Table S5a.** 2021

| F-value | | Pumpkin | | Barley | |
| --- | --- | --- | --- | --- | --- |
|  |  | Mono | Strip | Mono | Strip |
| Pumpkin | Mono |  |  |  |  |
|  | Strip | **3.64** |  |  |  |
| Barley | Mono | **10.39** | 1.40 |  |  |
|  | Strip | **5.82** | 1.21 | **4.23** |  |

| F-value | | Potato | | Grass | |
| --- | --- | --- | --- | --- | --- |
|  |  | Mono | Strip | Mono | Strip |
| Potato | Mono |  |  |  |  |
|  | Strip | **5.00** |  |  |  |
| Grass | Mono | 1.70 | 0.57 |  |  |
|  | Strip | **8.38** | 5.06 | 1.94 |  |

| F-value | | Cabbage | | Oat | |
| --- | --- | --- | --- | --- | --- |
|  |  | Mono | Strip | Mono | Strip |
| Cabbage | Mono |  |  |  |  |
|  | Strip | 1.47 |  |  |  |
| Oat | Mono | **2.08** | **3.52** |  |  |
|  | Strip | 2.06 | 2.07 | 2.92 |  |

**Table S5b.** 2022

| F-value | | Pumpkin | | Barley | |
| --- | --- | --- | --- | --- | --- |
|  |  | Mono | Strip | Mono | Strip |
| Pumpkin | Mono |  |  |  |  |
|  | Strip | 1.42 |  |  |  |
| Barley | Mono | **3.84** | **2.90** |  |  |
|  | Strip | 1.82 | 2.18 | 0.76 |  |

| F-value | | Potato | | Grass | |
| --- | --- | --- | --- | --- | --- |
|  |  | Mono | Strip | Mono | Strip |
| Potato | Mono |  |  |  |  |
|  | Strip | 1.42 |  |  |  |
| Grass | Mono | **7.14** | 3.26 |  |  |
|  | Strip | **4.01** | 3.24 | 3.58 |  |

| F-value | | Cabbage | | Oat | |
| --- | --- | --- | --- | --- | --- |
|  |  | Mono | Strip | Mono | Strip |
| Cabbage | Mono |  |  |  |  |
|  | Strip | **5.35** |  |  |  |
| Oat | Mono | **8.72** | **1.82** |  |  |
|  | Strip | 1.59 | 3.56 | **5.05** |  |

**Table S6. Effect of crop configuration on crop yield.** Yield results were retrieved from published and unpublished studies on effects of strip cropping on crop yield in similar locations, years, and crops as this study. Mean crop yield is presented in ton per hectare (t/ha). When known, standard deviations of mean crop yield are given (± SD). When a crop is indicated with NC (not collected) the crop yield was not collected due to an inconsistent sampling method (potato, 2020, Almere), crop failure (broccoli, 2020, Almere; celeriac, 2021, Almere), unavailable machine-harvest data (grass, Almere, 2020, 2021, 2022) and undocumented reasons (barley/beans, 2020, Valthermond). Unavailable data include cabbage (2019) and potato (2020) in Lelystad; and pumpkin (2020, 2021, 2022), barley (2020, 2021, 2022), oat (2021, 2022), potato (2021, 2022), grass (2021, 2022), and cabbage (2022) in Wageningen.

| Location | Year | Crops | Reference | Yield (t/ha) |  |
| --- | --- | --- | --- | --- | --- |
|  |  |  |  | Mono | Strip |
| Almere | 2020 | Beans | Juventia et al., 2024 | 2.80 | 3.45 |
|  |  | Broccoli |  | NC | NC |
|  |  | Celeriac |  | 54.44 | 69.57 |
|  |  | Grass |  | NC | NC |
|  |  | Oat |  | 6.05 | 5.08 |
|  |  | Onion |  | 41.06 | 42.14 |
|  |  | Parsnip |  | 27.50 | 35.36 |
|  |  | Potato |  | NC | NC |
|  | 2021 | Beans | Juventia et al., 2024 | 4.72 | 6.50 |
|  |  | Broccoli |  | 9.30 | 4.90 |
|  |  | Celeriac |  | NC | NC |
|  |  | Grass |  | NC | NC |
|  |  | Oat |  | 8.76 | 8.25 |
|  |  | Onion |  | 39.03 | 37.39 |
|  |  | Parsnip |  | 36.88 | 65.01 |
|  |  | Potato |  | 24.68 | 27.88 |
|  | 2022 | Beans | Juventia et al., 2024 | 4.12 | 4.36 |
|  |  | Broccoli |  | 1.12 | 1.59 |
|  |  | Celeriac |  | 45.38 | 51.45 |
|  |  | Grass |  | NC | NC |
|  |  | Oat |  | 6.27 | 6.70 |
|  |  | Onion |  | 57.89 | 48.62 |
|  |  | Parsnip |  | 20.6 | 21.09 |
|  |  | Potato |  | 31.08 | 31.60 |
| Lelystad | 2020 | Cabbage | Carillo-Reche et al., 2023 | 44.0 ± 12.8 | 46.3 ± 13.0 |
|  | 2021 | Cabbage | Carillo-Reche et al., 2023 | 85.6 ±18.2 | 75.9 ± 23.4 |
| Valthermond | 2020 | Barley | Unpublished data | 5.3 | 5.1 |
|  |  | Barley / Beans |  | NC | 5.1 |
|  |  | Potato |  | 20.9 | 20.5 |
|  | 2021 | Barley / Beans | Unpublished data | 3.1 | 3.4 |
|  |  | Grass / Clover |  | 8.3 | 7.9 |
|  |  | Potato |  | 23.7 | 20.7 |
| Wageningen | 2019 | Cabbage | Carillo-Reche et al., 2023 | 36.8 ± 15.8 | 34.6 ± 6.5 |
|  |  | Wheat | Ditzler et al., 2023 | 2.8 ± 0.2 | 2.1 ± 0.3 |
|  |  | Potato | Ditzler et al., 2023 | 29.3 ± 5.7 | 30.6 ± 8.8 |
|  | 2020 | Cabbage | Carillo-Reche et al., 2023 | 25.7 ± 9.3 | 19.3 ± 6.6 |
|  |  | Wheat | Ditzler et al., 2023 | 0.9 ± 0.2 | 1.0 ± 0.4 |
|  |  | Potato | Ditzler et al., 2023 | 23.4 ± 6.1 | 26.1 ± 8.9 |
|  | 2021 | Cabbage | Carillo-Reche et al., 2023 | 32.1 ± 7.1 | 27.9 ± 13.1 |

**Table S7. Sampling effort.** The total numbers of pitfall traps placed per location, year and crop. Rounds indicate the number of times pitfall traps were placed and were pooled within year series.

| Location | Year | Crops | Rounds | Pitfall traps per round | |
| --- | --- | --- | --- | --- | --- |
|  |  |  |  | Mono | Strip |
| Almere | 2020 | Beans | 3 | 4 | 4 |
|  |  | Broccoli |  | 4 | 4 |
|  |  | Celeriac |  | 4 | 4 |
|  |  | Grass |  | 4 | 3 |
|  |  | Oat |  | 4 | 4 |
|  |  | Onion |  | 4 | 4 |
|  |  | Parsnip |  | 4 | 4 |
|  |  | Potato |  | 4 | 4 |
|  | 2021 | Beans | 4 | 4 | 4 |
|  |  | Broccoli |  | 4 | 4 |
|  |  | Celeriac |  | 4 | 4 |
|  |  | Grass |  | 3 | 4 |
|  |  | Oat |  | 4 | 4 |
|  |  | Onion |  | 4 | 4 |
|  |  | Parsnip |  | 4 | 4 |
|  |  | Potato |  | 4 | 4 |
|  | 2022 | Beans | 5 | 4 | 4 |
|  |  | Broccoli |  | 4 | 3 |
|  |  | Celeriac |  | 3 | 2 |
|  |  | Grass |  | 4 | 4 |
|  |  | Oat |  | 4 | 2 |
|  |  | Onion |  | 3 | 4 |
|  |  | Parsnip |  | 4 | 4 |
|  |  | Potato | 4 | 4 | 4 |
| Lelystad | 2019 | Cabbage | 5 | 4 | 3 |
|  | 2020 | Cabbage | 7 | 4 | 3 |
|  |  | Potato | 2 | 4 | 2 |
|  | 2021 | Cabbage | 6 | 4 | 3 |
| Valthermond | 2020 | Barley | 7 | 6 | 6 |
|  |  | Barley / Beans |  | 6 | 6 |
|  |  | Potato |  | 6 | 6 |
|  | 2021 | Barley / Beans | 8 | 6 | 6 |
|  |  | Grass / Clover |  | 6 | 6 |
|  |  | Potato |  | 6 | 6 |
| Wageningen | 2019 | Cabbage | 4 | 6 | 2 |
|  |  | Wheat | 2 | 6 | 2 |
|  |  | Potato | 3 | 6 | 6 |
|  | 2020 | Cabbage | 8 | 6 | 2 |
|  |  | Wheat | 3 | 5 | 2 |
|  |  | Potato | 1 | 6 | 5 |
|  |  | Pumpkin | 2 | 6 | 4 |
|  |  | Barley |  | 6 | 2 |
|  | 2021 | Cabbage | 7 | 6 | 3 |
|  |  | Oat |  | 6 | 3 |
|  |  | Potato | 2 | 6 | 3 |
|  |  | Grass |  | 6 | 3 |
|  |  | Pumpkin |  | 6 | 3 |
|  |  | Barley |  | 6 | 3 |
|  | 2022 | Cabbage | 2 | 6 | 3 |
|  |  | Oat |  | 5 | 3 |
|  |  | Potato |  | 6 | 3 |
|  |  | Grass |  | 6 | 3 |
|  |  | Pumpkin |  | 6 | 3 |
|  |  | Barley |  | 6 | 3 |

**Table S8.** Plant species composition of flower strips adjacent to the strip cropping fields in Lelystad and Valthermond (see location in Fig. S10).

|  | **Species** |
| --- | --- |
| 1 | *Achillea millefolium* (common yarrow) |
| 2 | *Anethum graveolens* (dill) |
| 3 | *Cichorium intybus* (common chichory) |
| 4 | *Foeniculum vulgare* (fennel) |
| 5 | *Pastinaca sativa* (parsnip) |
| 6 | *Angelica sylvestris* (wild angelica) |
| 7 | *Anthriscus sylvestris* (cow parsley) |
| 8 | *Fagopyrum esculentum* (buckwheat) |
| 9 | *Papaver Rhoeas* (common poppy red wild form) |
| 10 | *Chrysanthemum segetum* (corn marigold) |
| 11 | *Ammi majus* (greater ammi) |
| 12 | *Centaurea cyanus* (cornflower, single flowered wild form) |
